# Mechanical principles governing the shapes of dendritic spines

**DOI:** 10.1101/2020.09.09.290650

**Authors:** H. Alimohamadi, M.K. Bell, S. Halpain, P. Rangamani

**Affiliations:** Department of Mechanical and Aerospace Engineering, University of California San Diego, CA 92093, USA; Sanford Consortium for Regenerative Medicine, La Jolla 92037, USA; Section of Neurobiology, Division of Biological Sciences, University of California San Diego, La Jolla 92037, USA

**Keywords:** Lipid bilayer, Dendritic spines, Membrane-actin interactions, Deviatoric curvature

## Abstract

Dendritic spines are small, bulbous protrusions along the dendrites of neurons and are sites of excitatory postsynaptic activity. The morphology of spines has been implicated in their function in synaptic plasticity and their shapes have been well-characterized, but the potential mechanics underlying their shape development and maintenance have not yet been fully understood. In this work, we explore the mechanical principles that could underlie specific shapes using a minimal biophysical model of membrane-actin interactions. Using this model, we first identify the possible force regimes that give rise to the classic spine shapes – stubby, filopodia, thin, and mushroom-shaped spines. We also use this model to investigate how the spine neck might be stabilized using periodic rings of actin or associated proteins. Finally, we use this model to predict that the cooperation between force generation and ring structures can regulate the energy landscape of spine shapes across a wide range of tensions. Thus, our study provides insights into how mechanical aspects of actin-mediated force generation and tension can play critical roles in spine shape maintenance.

## 1 Introduction

Dendritic spines are small, bulbous protrusions along the dendrites of neurons that occur at postsynaptic glutamatergic synapses [1–3]. They respond to a glutamate release by orchestrating a series of biochemical and biophysical events that span multiple spatial and temporal scales [4–6]. Spine morphology is tightly coupled to synaptic function, with larger spines tending to represent stronger synapses [7, 8] due to their greater surface expression of functional glutamate receptors. Synaptic activity regulates spine shape and volume. For example, several forms of physiological synaptic plasticity, such as long-term potentiation (LTP) and long-term depression (LTD) are associated with spine enlargement and spine shrinkage, respectively [9–11]. Although average spine volume is approximately 0.1 femtoliter, the shape and volume of dendritic spines are highly variable, depending both on the developmental stage and a combination of genetic and environmental factors, including the prior history of activity [12–15]. Moreover, spine morphology is highly dynamic on the scale of seconds to minutes, due to a dynamic actin-based cytoskeleton [2, 16].

Despite their broad range of morphological features and highly dynamic nature, dendritic spines can be classified into four broad categories. Spines in the mature nervous system are typically classified as being stubby, thin, or mushroom-shaped [17, 18] (Fig. 1A). These categories of spines can be identified in electron micrographs as postsynaptic structures connected to presynaptic nerve terminals. Stubby spines are short and wide, and lack a discernible neck. Such spines appear early during synaptogenesis and may represent an emerging spine, but they also might result from spine shrinkage driven by physiological or pathological conditions (Fig.1A) [4, 12, 19].

**Figure 1:**
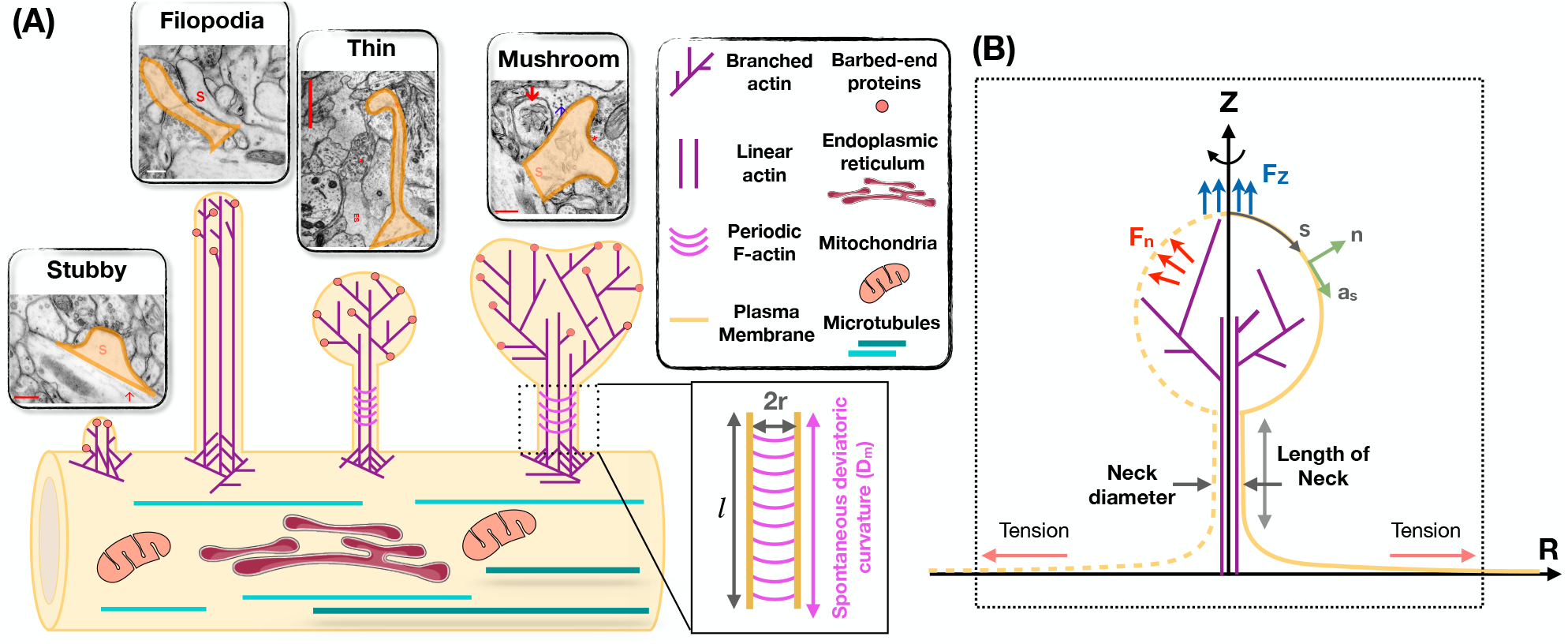
Modeling of forces relevant to spine shape. (A) Schematic depiction of different shape categories of dendritic spines (Reprinted with permission from SynapseWeb, Kristen M. Harris, PI, http://synapseweb.clm.utexas.edu/). The inset shows a schematic of a tubular neck with a radius r and a spontaneous deviatoric curvature D_m_ along the total neck length *l*. (B) The surface parametrization of the membrane geometry in axisymmetric coordinates. s is the arclength, **n** is the unit normal vector to the membrane surface, and **a**_*s*_ is the unit tangent vector in the direction of arclength. We assume that the actin filaments can apply axial (F_z_) or normal (F_n_) forces to the membrane surface. We assume that there is a large membrane reservoir with a fixed area, and we focused on the local region of the membrane under tension *λ*, as indicated by the dotted box.

The adult mammalian brain is dominated by either thin or mushroom-shaped spines. Thin spines have a long thin neck that is connected to a small bulbous head (Fig. 1A) [12]. Within the head is the postsynaptic density (PSD), an area just beneath the synaptic plasma membrane containing a high concentration of glutamate receptors, scaffolding molecules, and other proteins essential for postsynaptic function. Thin spines have flexible structures that allow them to adapt their morphology based on different levels of synaptic activity [20, 21]. It has been proposed that thin spines are “learning spines,” because they display a high capacity for expansion and strengthening via insertion of new AMPA-type glutamate receptors into the PSD, which is the key basis for synapse strengthening [20–24]. Compared to thin spines, mushroom-shaped spines have a shorter neck and a greatly expanded head (Fig. 1A) [12]. Mature mushroom-shaped spines are more likely to be stable for months to years [20, 21, 24–26], with slower turnover, and are associated with strong synapse functionality, as they contain on average higher concentrations of AMPA-type glutamate receptors. Such spines have therefore been called “memory spines”, in the sense that their potentiated strength reflects a history of high activity and thus “memory” storage, yet their capacity for further potentiation may be near saturation [22–24, 27–29]. Table 1 provides the reported dimensions for different shape categories of dendritic spines observed in hippocampal neurons [18, 30–33].

**Table 1:** Dimensions of different spine shapes compiled from the literature.

|  | Stubby [18, 32] | Filopodia [30, 31] | Thin [18, 32, 33] | Mushroom [18, 33] |
| --- | --- | --- | --- | --- |
| Total length ($L$) ($\mu\text{m}$) | $0.44 \pm 0.15$ | 2-20 | $0.98 \pm 0.42$ | $1.5 \pm 0.25$ |
| Length of neck ($l$) ($\mu\text{m}$) | – | – | $0.51 \pm 0.34$ | $0.43 \pm 0.21$ |
| Neck diameter ($2r$) ($\mu\text{m}$) | $0.32 \pm 0.13$ | $< 0.3$ | $0.1 \pm 0.03$ | $0.2 \pm 0.07$ |
| Total volume ($\mu\text{m}^3$) | $0.03 \pm 0.01$ | – | $0.04 \pm 0.02$ | $0.29 \pm 0.13$ |
| Volume of head ($V$) ($\mu\text{m}^3$) | – | – | $0.03 \pm 0.15$ | $0.27 \pm 0.13$ |
| Total surface area ($\mu\text{m}^2$) | $0.45 \pm 0.14$ | – | $0.59 \pm 0.29$ | $2.7 \pm 0.93$ |
| Surface area of head ($\mu\text{m}^2$) | – | – | $0.4 \pm 0.15$ | $2.4 \pm 0.92$ |
| Surface area of PSD ($\mu\text{m}^2$) | $0.07 \pm 0.02$ | – | $0.05 \pm 0.02$ | $0.3 \pm 0.1$ |
| Surface area of PSD/head | – | – | $0.1 \pm 0.06$ | $0.18 \pm 0.15$ |

In addition to synapse-bearing spines, the fourth category of spine-like protrusions is dendritic filopodia. These are commonly observed during early development, and are thought to facilitate the pairing of presynaptic and postsynaptic glutamatergic sites during synaptogenesis by spatially scanning the neuropil volume for a partner axon [19, 34–36]. Thus, a fraction of these “protospines” become synapse-bearing spines if they come into contact with and are stabilized in partnership with presynaptic nerve terminals [34, 37]. Filopodia are long (>2 *μ*m) and thin (< 0.3 *μ*m diameter) protrusions that lack a bulbous head (Fig. 1A) [30].

Because the size and shape of functional subcellular domains are closely tied to the mechanics of actin-membrane interactions [18, 30, 31], a more complete understanding of dendritic spine dynamics, development, and function would benefit from biophysical models that address the underlying mechanical aspects. We have therefore begun to build a computational model of spines that incorporates both membrane forces and actin-based forces, and their interaction. This model is based on published experimental observations in dendritic spines, non-neuronal cells, and biochemical experiments. The goal of this model is to inform our understanding of the development of spines and the plasticity of their structure under different physiological scenarios.

Currently, there are hundreds of studies that address various aspects of the regulation of dendritic spine size and shape. In building our model, we have chosen to focus on several key observations, as follows.

1. **Actin enrichment in spines:** Dendritic spines are enriched in filamentous actin, which, along with scaffolding molecules, establish spine architecture [38–40]. Membrane-actin interactions associated with spine enlargement and shrinkage during plasticity can be modeled at the single filament level using the elastic Brownian ratchet and the net force acting on the membrane due to actin remodeling can be represented as work done by actin to deform the membrane [41–43].
2. **Different subpopulations of actin:** There appear to be distinct subpopulations of F-actin in dendritic spines, and spine actin can be thought of as an independent network with interconnected nodes [44]. The spine head typically consists of short, cross-linked filaments; branched filaments have been observed in the spine head [39, 45, 46]. The spine neck was initially thought to contain long filaments [47–50], but current evidence has suggested the presence of short, branched filaments [45]. Additionally, recent high resolution imaging techniques have shown that there are likely periodic F-actin structures along the neck region of dendritic spines [51, 52]. These periodic F-actin structures are very stable and in contrast to long and branched filaments, resist depolymerization [52].
3. **Roles for actin binding proteins:** Actin dynamics in spines are tightly regulated by dozens of various actin binding proteins, some of which must also interact directly or indirectly with the spine plasma membrane [53–55]. First, the turnover of filaments themselves can drive forces against the membrane that regulate the expansion, maintenance, or shrinkage of spine compartments [56]. The key factors that govern this balance are (a) the rate of polymerization, which is regulated by actin nucleating factors such as formins and the Arp2/3 complex [39]; (b) the rate of depolymerization, which is regulated by actin severing factors such as cofilin and gelsolin [57]; (c) the number of free barbed ends, which is regulated via actin severing activity and the activity of barbed end capping proteins [58]; and (d) the concentration of available actin monomer, which is dependent upon the G-actin concentration and also the activities of profilin, which delivers ATP-bound G-actin to the above actin nucleators, and regulators such as N-WASP, which controls Arp2/3 activity [58, 59]. In addition, several proteins that crosslink or stabilize actin filaments, such as cortactin [60], spectrin [61], or drebrin [62] are known to regulate spine shape and separately, myosin motors can affect spine shape either directly by creating contractile forces, or indirectly by regulating the transport of cargo into and out of the spine [58, 63]. Finally, an important role for calcium/calmodulin-dependent protein kinase in structural plasticity of spines has been demonstrated through its ability not only to transduce calcium signals, but also to regulate actin directly through the direct binding of F-actin via its *β*-subunit [64–67].
4. **Membrane mechanics:** All cells regulate their shape by coordinating the properties of the cytoskeleton with that of the plasma membrane. Proteins such as MARCKS that interact directly with both F-actin and the lipid bilayer can strongly influence spine shape [68]. Membrane curvature is especially important in spines and represents a specific mechanical force that is regulated by specific proteins, as well as lipid composition. Bin/Amphiphysin/Rvs (BAR)-domain containing proteins assemble on the membrane to produce anisotropic curvature and promote tubulation. Studies have demonstrated critical roles for specific BAR-domain proteins in dendritic spines. Recently, the role of membrane mechanics has been elucidated in the initiation of dendritic spines [69]. A series of studies showed that dendritic spines can be initiated by membrane bending due to protein patches containing BAR domains such as I-BAR and F-BAR proteins [70–73]. These proteins are known to polymerize on the membrane [74–77], induce anisotropic curvature [78–80], and promote tubulation [76, 81–84].

The above findings suggest that membrane bending and actin-membrane interactions are major determinants of spine morphology. Recent studies have modeled the role of either membrane mechanics alone [43] or actin dynamics alone in spines [85], but the interaction between the two has not yet been addressed. Here, we present a general theoretical model that relates membrane bending and actin-mediated forces to spine morphology. Using this model, we investigate the mechanical landscape of the different shapes of spines and map the relationships among actin-mediated force generation, membrane elasticity, and curvature induced by periodic ring structures and proteins such as BAR domains.

## 2 Model development

### 2.1 Assumptions

- We treat the lipid bilayer as a continuous thin elastic shell, assuming that the membrane thickness is negligible compared to the radii of membrane curvature [86, 87]. This allows us to model the bending energy of the membrane using the modified version of the Helfrich–Canham energy, including the effect of spatially varying deviatoric curvature to represent the induced anisotropic curvatures by periodic F-actin rings and other ring-shaped structures [81, 82, 87–90].
- We assume that the membrane is locally inextensible, since the stretching modulus of the lipid bilayer is an order of magnitude larger than the membrane bending modulus [91]. We implemented this constraint using a Lagrange multiplier, which can be interpreted as the membrane tension [92–94]. We note that this membrane tension, in this study, is better interpreted as the cortical tension including the effective contribution of both the membrane in-plane stresses and membrane-cytoskeleton interactions [95, 96].
- We assume that the time scales of mechanical forces are much faster than other events in dendritic spines, allowing us to assume mechanical equilibrium and neglect inertia [43, 92]. This assumption is justified by the fact that the timescale of the equilibration of the mechanical forces is much smaller than the timescale of actin polymerization in dendritic spines [97].
- We assume that the force exerted by the actin cytoskeleton can be represented as work done on the membrane and do not include the molecular details of the actin network [43, 96, 98–100]. Additionally, we assume that the periodic ring shaped structures of actin and related proteins such as *β*II spectrin and BAR-domain proteins can be represented using an anisotropic spontaneous curvature [80–82, 89].
- For ease of computation we assume that the geometry of a dendritic spine is rotationally symmetric (see Fig. 1B) [43]. This assumption allows us to parametrize the whole surface by a single parameter, arclength.

### 2.2 Mechanical force balance

In this section we present a concise derivation of the governing mathematical shape equations for the shape of dendritic spines at mechanical equilibrium. The complete derivation with details is given in [92, 98, 101, 102]. The total free energy of the system (*E*) includes the elastic storage energy of the membrane (*E*_elastic_), and the work done by the applied forces due to actin filaments (*W*_force_) [90, 98, 103, 104] is given by

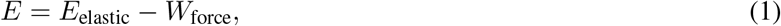

where

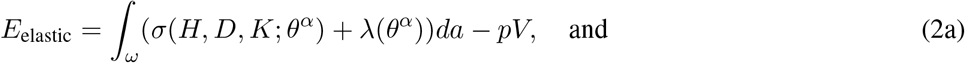

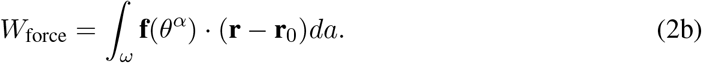

Here, *ω* is the total membrane surface area, *σ* is the bending energy density per unit area, *θ*^*α*^ denotes the surface coordinate where *α* ∈ {1, 2}, *H* is the mean curvature of the surface, *D* is the curvature deviator, *K* is the Gaussian curvature of the surface, *λ* is the tension field and represents the Lagrange multiplier associated with the local area constraint, *p* is the transmembrane pressure and represents the Lagrange multiplier associated with the volume constraint, *V* is the enclosed volume, **f** is the applied force per unit area, **r** is the position vector in the current configuration, and **r**_0_ is the position vector in the reference frame. To model the energy density *σ* in Eq. 2a, we used the modified version of Helfrich energy including the effects of induced anisotropic curvature by periodic F-actin structures and BAR domain proteins [78, 81, 82, 87, 88, 90, 105], given as

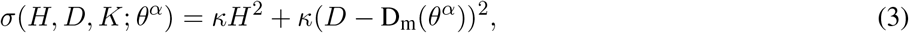

where *κ* is a constant representing the bending moduli and D_m_ is the spontaneous (intrinsic) deviatoric curvature which can be spatially heterogeneous along the membrane surface [78, 81, 82]. It should be mentioned that in Eq. 3, we assumed that periodic rings can only induce anisotropic curvature and we set the isotropic curvature (spontaneous curvature) to be zero throughout this study. Substituting Eqs. 2a, 2b, and 3 into Eq. 1 gives

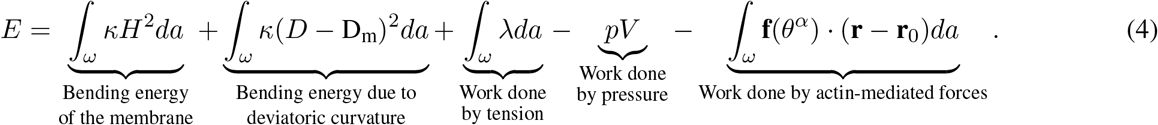

Minimization of the energy (Eq. 4) using the variational approach results in the governing shape equation (Eq. S6) and the incompressibility condition (Eq.S7) for a heterogeneous membrane. The complete equations are presented in the supplementary material along with the complete notation in Table S1.

### 2.3 Numerical implementation

In axisymmetric coordinates, the membrane shape equation (Eq. S6) and the incompressibility condition (Eq.S7) simplify to a coupled system of first order differential equations (Eq. S24). In order to solve this system of equations along with the prescribed boundary conditions (Eq. S25), we used ‘bvp4c,’ a boundary value problem solver in MATLAB. In all our simulations, we assume that the total area of the membrane is conserved and we also fixed the bending modulus to be *κ* = 0.18 pN*·µ*m based on previous models for spines [106, 107]. We also set the transmembrane pressure to zero (*p* = 0) to focus only on the mechanism of membrane-actin interactions in governing the shapes of dendritic spines.

## 3 Results

Using the model described above, we conducted simulations for different mechanical parameters with the goal of identifying the range of forces, the associated heterogeneities, and the protein-induced and cytoskeleton-induced anisotropic curvatures that could result in shapes and sizes of spines corresponding to those observed experimentally (Table 1). Specifically, we sought to recreate the filopodial, stubby, thin, and mushroom-shaped spines as shown in Fig. 1. We must emphasize that all the shapes are equilibrium shapes, and our model does not provide insight into dynamic transitions from one shape to another. Our simulation results are described below. In these data, we emphasize the relationships among different mechanical parameters to obtain the desired shapes, and give specific values for mechanical parameters that result in sizes as listed in Table 1. These provide some realistic magnitudes for forces present at various locations within the compact spine volume.

### 3.1 Localized axial forces along the membrane are sufficient for the formation of stubby and filopodial shaped spines

We begin with an analysis of the force-shape relationship of stubby spines. We assumed that actin filaments exert axial forces in the nascent PSD area, which is a small fraction of the membrane surface area (Table 1). This heterogeneous force distribution along the membrane was implemented using a hyperbolic tangent function (Eq. S26). We observed that the relationship between the magnitude of the forces and the length of the stubby spines depends on the value of tension. To map this relationship, we performed the simulation for (*i*) a fixed height (L = 0.44 *µ*m) and a wide range of tensions (Fig. 2A) and (*ii*) a fixed tension (e.g., *λ* = 10 pN*/µ*m) and different heights of the stubby spine (Fig. 2B). As shown in previous studies [108, 109], for a small membrane deformation, such as a stubby spine, the axial force is linearly proportional to both tension and the height of the stubby spine (Figs. 2A & B) [108, 109]. Thus, from a mechanical standpoint, the stubby spine shape is accessible for a wide range of forces and tensions in the physiological range. If we seek to match a specific height of the stubby spine as noted from experimental observations, our simulations showed that an axial force of F_z_ = 7.5 pN is required to form a stubby spine of the length of L = 0.44 *µ*m (Table 1) when the tension is *λ* = 10 pN*/µ*m (Fig. 2C).

**Figure 2:**
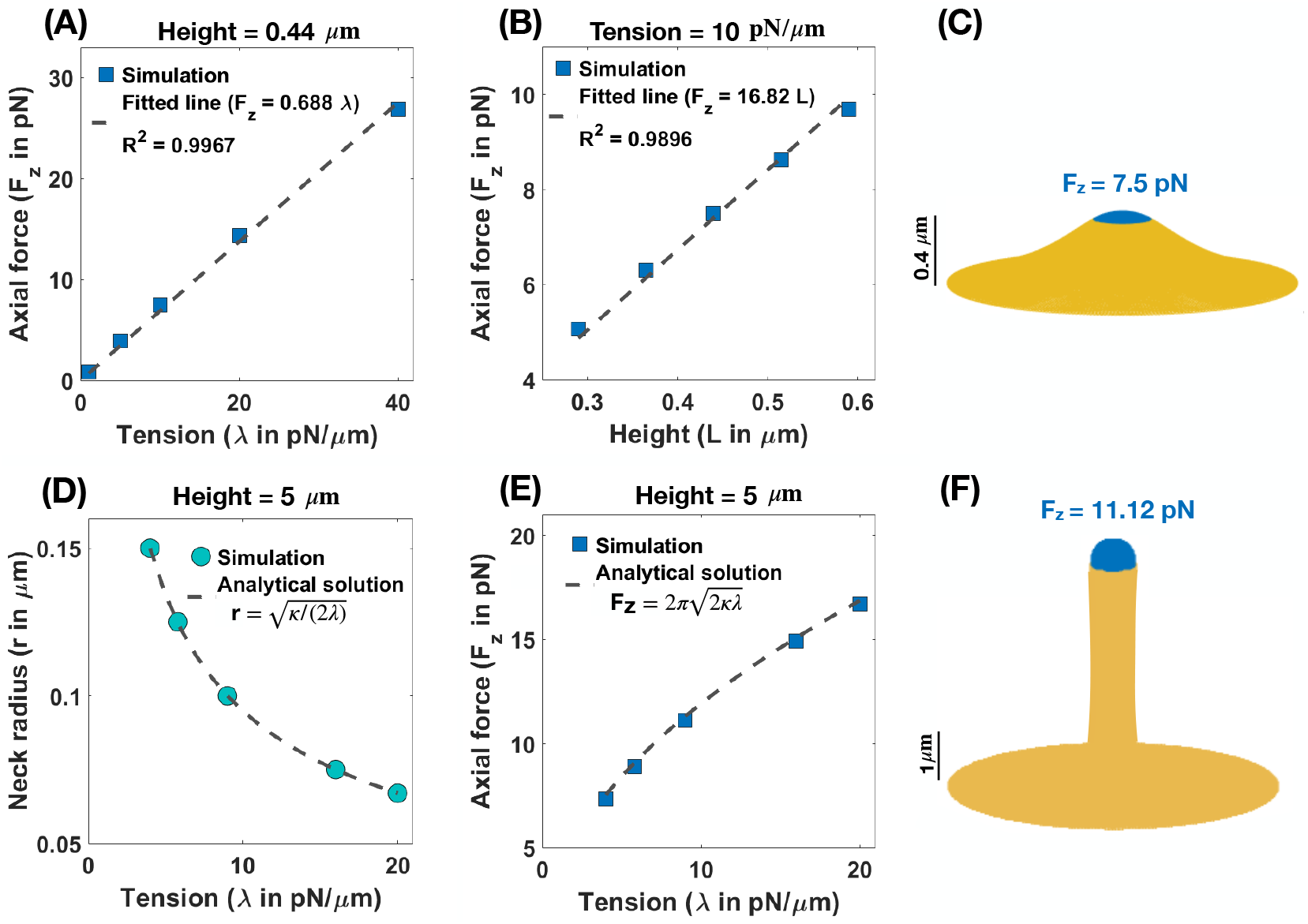
Formation of stubby and filopodia shaped spines with a localized axial force. (A) Linear relationship between the magnitude of axial force and tension in a small stubby-shaped membrane deformation [108, 109]. The dashed line is the fitted curve (F_z_ = 0.688*λ*) with R^2^ = 0.9967. (B) Linear relationship between the magnitude of axial force and the height of the stubby spine for a fixed tension [108, 109]. The dashed line is the fitted curve (F_z_ = 16.82L) with R^2^ = 0.9896. (C) A stubby-shaped spine with a total length L = 0.44 *µ*m is formed with F_*z*_ = 7.5 pN applied along the blue area (*λ* =10 pN*/µ*m). (D) Neck radius of a filopodium as a function of tension 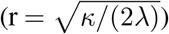 [108]. (E) The magnitude of axial force needed to form a filopodium as a function of tension 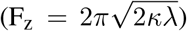 [108]. (F) A filopodium-shaped protrusion with a total length L = 5 *µ*m and neck radius r = 0.2 *µ*m is formed with F_*z*_ = 11.2 pN applied along the spherical cap of the filopodium, which is shown in blue (*λ* = 9 pN*/µ*m).

Next, we investigated the role of forces in the formation of long spines that resembled filopodia. As expected, we found that the formation of a long filopodium follows well-established results for tube formation in membranes [108]. Ignoring the spherical cap, a filopodium is a tubular membrane and its equilibrium radius (r) depends on the tension and bending rigidity of the membrane as 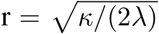 (Fig. 2D) [108]. The axial force F required to maintain the tubule with radius r and height L is given as 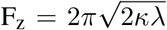 (Fig. 2E) [108]. Results from our simulations are consistent with these theoretical relationships. Specifically, for a filopodial-shaped protrusion of L = 5 *µ*m with a neck radius of r = 0.1 *µ*m, the required force was F_z_ = 11.12 pN for a tension of *λ* = *κ/*(2r^2^) = 9 pN*/µ*m (Fig. 2F) and the magnitude of this axial force is independent of the length of the protrusion (Fig. S3A).

### 3.2 Normal forces along the membrane support the formation of thin shaped spines

We next investigated the nature of forces that could be associated with the formation of thin-shaped spines. Because thin-shaped spines have a bulbous head, axial forces such as those used in Fig. 2 are insufficient to generate the spherical shape of the head. Since spherical shapes can be obtained by a normal force acting locally on the head region, we repeated the simulation in Fig. 2 but now included a localized uniform normal force density along the area of the spine head (*A*_force_ = *A*_spine head_). It is possible that such normal forces result from the dense actin meshwork in the spine heads [43, 110]. We estimated the forces required to generate a spherical head by assuming that a thin spine is ideally a sphere with radius R which is connected to a cylinder with radius r and height *l* (Fig. S1B). If a uniform normal force density, f_n_, is applied all along the sphere, then, ignoring the interface between the sphere and the cylinder, the total energy of the system can be written as

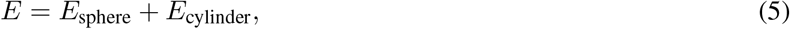

where *E*_sphere_ = (*κ/R*^2^ + *λ*)4*πR*^2^ − (4*π/*3)*R*^3^f_n_ and 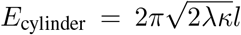 (see section 1.5.2 in the supplementary material). Minimizing the total energy of the system with respect to R by taking ∂*E/*∂*R* = 0, we obtain the equilibrium normal force density as f_n_ = 2*λ/R*. This resembles the Young-Laplace equation where normally pressure (normal force density) is a global parameter; in this case, f_n_ is a local normal force density. To find the total magnitude of normal force, we need to multiply the force density by the area of the sphere, which produces F_n_ = 8*πRλ*. In our simulation, we prescribe the area of the applied force and can rewrite the force as

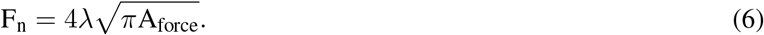

In order to generate thin-shaped spines, we first fixed the neck diameter based on the magnitude of tension 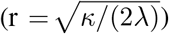 as shown in Fig. 2D. Similar to filopodia, in thin spines, the radius of the neck is related to the tension and the bending rigidity, given by 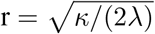 [108] (Fig. 3A). This relationship suggests that in order to have a thin spine with a neck radius between 0.035 *µ*m *<* r *<* 0.065 *µ*m (given range in Table 1), the tension can vary between 20 pN*/µ*m *< λ <* 80 pN*/µ*m. Based on Eq. 6, the magnitude of the normal force linearly depends on the tension, while it varies as the square root of the area of applied force.

**Figure 3:**
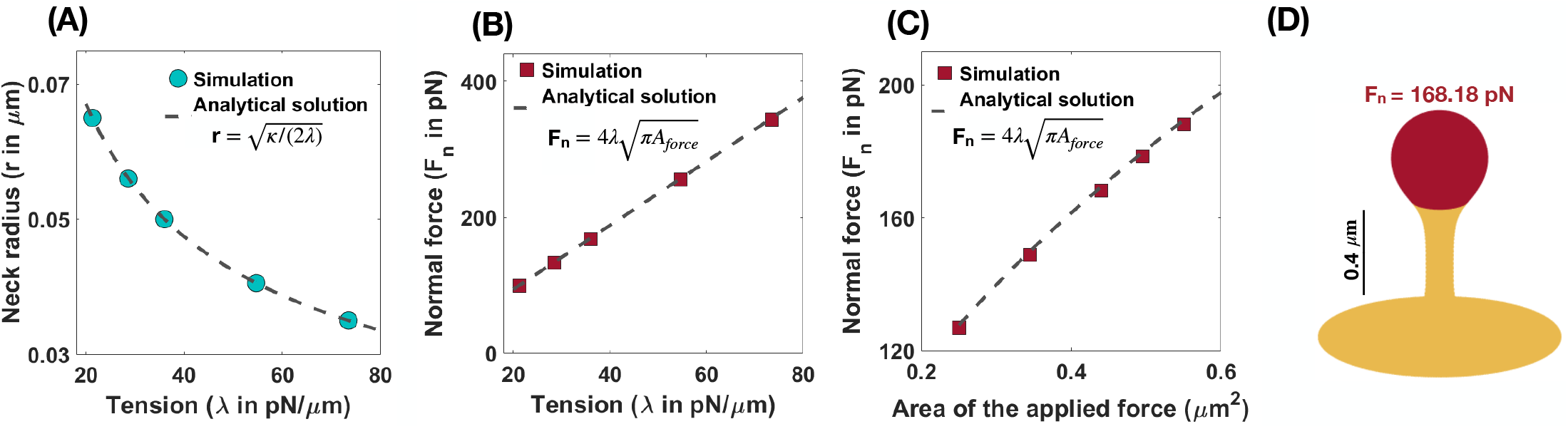
Formation of thin-shaped spines with localized normal force along the spine head. (A) Neck radius of a thin-shaped spine as a function of tension 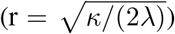 [108]. (B) Linear relationship between the magnitude of normal force needed to form a thin-shaped spine and the tension. Here the area of the applied force is set at *A*_force_ = 0.44 *µ*m^2^. The red squares represent the results obtained from simulation and the dashed line is the derived analytical solution 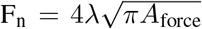, Eq. 6). (C) The magnitude of a normal force needed to form a thin-shaped spine as a function of the area of the spine head. The tension is set at *λ* =36 pN*/µ*m. The red squares represent the results obtained from our simulations and the dashed line is the derived analytical solution 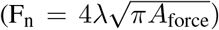), Eq. 6. (D) A thin-shaped spine with a total length L = 0.98 *µ*m, neck radius r = 0.05 *µ*m, and head volume V = 0.033 *µ*m^3^ is formed with F_n_ = 168.18 pN applied along the head of spine which is shown in red (*λ* =36 pN*/µ*m and *A*_force_ = 0.44 *µ*m^2^).

In Fig. 3B, we plotted the magnitude of the normal force as a function of tension obtained from numerical solutions (red squares) versus the analytical expression given in Eq. 6 (dotted line) for fixed *A*_force_ = 0.44 *µ*m^2^. We found a good agreement between the analytical solution and the results obtained from simulation such that by changing tension between 20 pN*/µ*m *< λ <* 80 pN*/µ*m, the magnitude of the normal force required to form a thin-shaped spine varies in a large range between 150 pN *<* F_n_ *<* 400 pN (Fig. 3B). To further validate our numerical results, we plotted the magnitude of the normal force as a function of the area of the applied force (*A*_force_) obtained from numerical solution (red squares) versus the analytical expression given in Eq. 6 (dotted line) for a fixed tension, *λ* = 36 pN*/µ*m (Fig. 3C). We observed a good agreement between the analytical solution and the numerical results where by increasing the area of the applied force from *A*_force_ = 0.25 *µ*m^2^ to *A*_force_ = 0.55 *µ*m^2^, the magnitude of the normal applied force needed to form a thin spine increases from F_n_ ∼ 120 pN to F_n_ ∼ 200 pN (Fig. 3C).

Thus, to form a thin spine with an average neck diameter of r = 0.05 *µ*m (see Table 1), we set our tension to be *λ* =36 pN*/µ*m 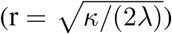. Based on our calculation for *λ* =36 pN*/µ*m and *A*_force_ = 0.44 *µ*m^2^ (average area of the spine head in Table 1), a total normal force of F_n_ = 168.18 pN (applied along the red area) is required to form a thin spine with a total length L = 0.98 *µ*m, a neck radius r = 0.05 *µ*m, and a head volume V = 0.033 *µ*m^3^ (Fig. 3D). Also, in Fig. S3B, we show that the magnitude of the normal force needed to form a thin spine is independent of the height of the spine.

### 3.3 Non-uniform normal force distributions can result in mushroom-shaped spines

We next asked if changes to the force distributions could result in mushroom-shaped spines. We hypothesized that one possible way is to have a heterogeneous force distribution along the spine head and the PSD area. To understand how non-uniform distributions of normal forces can characterize the morphology of mushroom spines, we performed simulations assuming that the normal force applied along the PSD area is different from the normal force applied along the rest of the spine head (Fig. 4A).

**Figure 4:**
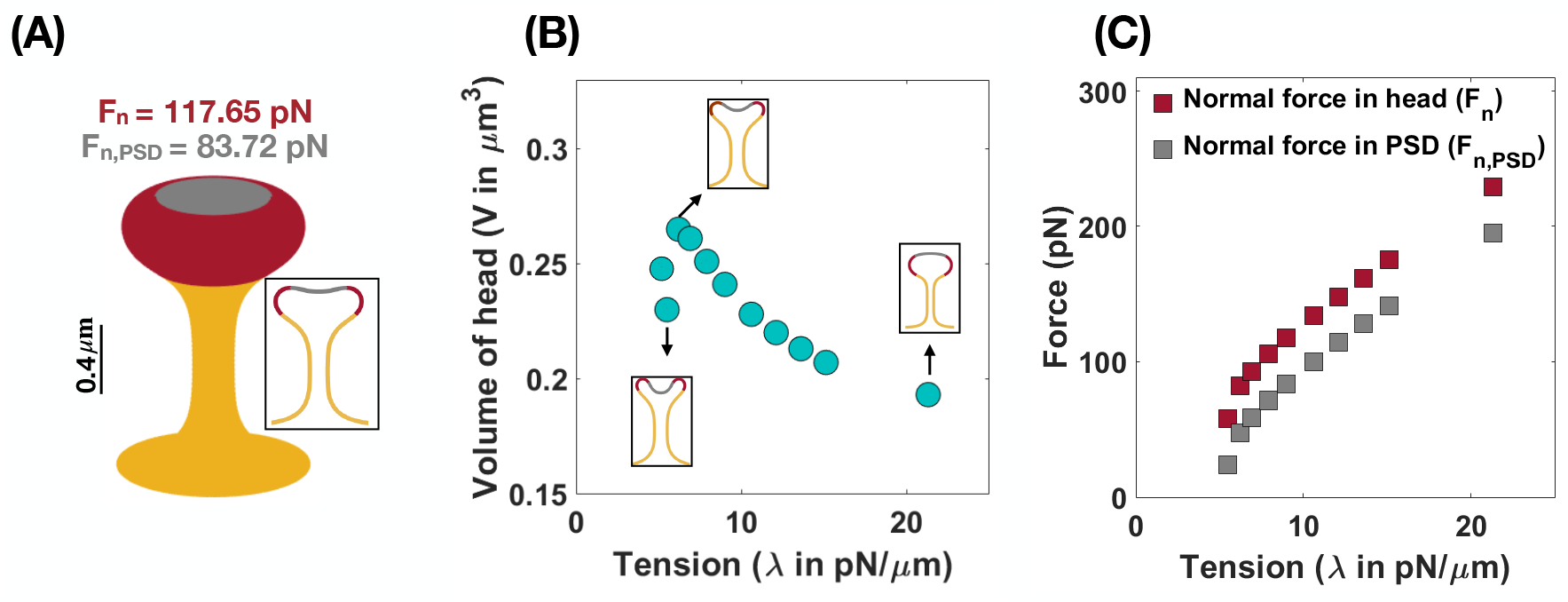
Formation of mushroom-shaped spines with localized normal forces along the spine head and PSD. (A) A mushroom-shaped spine with a total length L = 1.51 *µ*m, neck radius r = 0.1 *µ*m, head volume V = 0.25 *µ*m^3^, and area of PSD/area of head = 0.2 is formed with F_n_ = 117.65 pN applied along the head of spine (red domain) and F_n,PSD_ = 83.72 pN applied along the PSD (gray domain) (*λ* =9 pN*/µ*m). (B) The nonmonotonic behavior of the volume of a mushroom-shaped spine head when increasing tension. Three different shapes of mushroom-shaped spines are shown for low, intermediate, and high tensions. With increasing magnitude of tension, the mushroom-shaped spine head flattens. (C) The magnitude of normal forces in the spine head (red squares) and in PSD (gray squares) increases with increasing tension.

In the case of mushroom-shaped spines, we have multiple geometric parameters to consider – (a) head volume, (b) area fraction of the PSD, and (c) tension. For example, to form a mushroom-shaped spine with a total length L = 1.51 *µ*m, head volume V = 0.25 *µ*m^3^, and area of PSD/ area of head ratio = 0.2 (see Table 1), normal forces of F_n_ = 117.65 pN and F_n,PSD_ = 83.72 pN are required along the spine head (red region) and the PSD area (gray region), respectively (Fig. 4A). The value of tension was set to *λ* = 9 pN*/µ*m to obtain a neck radius of about r ≈ 0.1 *µ*m (see Table 1 and Fig. S4). The magnitude of these forces is independent of the height of the spine (Fig. S3C).

We observed that the morphology of the spine head changes with varying magnitude of tension; the spine head flattens for large tensions (Fig. 4B). This is consistent with previous studies that have investigated membrane shape at high tensions, e.g., the membrane remains almost flat during vesicle budding [111, 100], or in the case of a red blood cell, the biconcave cell flattens to a pancake shape [112, 96]. To further investigate how a change in the morphology of the spine head can affect the volume of the head, we plotted the volume of the head (V) as a function of tension (Fig. 4B). We found that the head volume is a non-monotonic function of tension; as tension increases, the volume of the spine head increases and then decreases (Fig. 4B).

This is because initially when increasing tension from low to intermediate values the head flattens and the volume of the head increases. However, for high tensions, the shrinkage of the head becomes dominant and as a result the volume decreases (Fig. 4B). Consistent with these observations, a larger normal force is required to bend a stiffer membrane and form a mushroom-shaped spine (Fig. 4C). For example, based on our calculation, when increasing tension from *λ* = 5 pN*/µ*m to *λ* = 20 pN*/µ*m, the normal forces in the spine head and PSD area increase by almost 150 pN (Fig. 4C).

To study how the ratio of PSD area to the total area of the spine head affects the magnitude of normal forces, we performed simulations for a range of area of PSD/area of head ratios (Fig. S5). Our results show that with increasing area of PSD/area of head ratio, larger normal forces both in the spine head and the PSD region are required (Fig. S5A). Additionally, increasing the ratio of the PSD area to the total area of the head results in the flattening of the spine head with a larger volume (Fig. S5B). Thus, mushroom-shaped spines can be formed from a multitude of mechanical pathways – heterogeneous forces in the spine head, balancing tension and force distributions, and using different area localizations of the forces.

### 3.4 Induced spontaneous deviatoric curvature by periodic F-actins structures and BAR domain proteins can generate characteristic dendritic spine necks

Recently, super-resolution microscopy methods have revealed the presence of ubiquitous actin ring structures along spine necks [51, 52]. It has been suggested that these ring-like structures and BAR-domain proteins can together support the tubular shape of dendritic spines [38, 113]. To understand how periodic F-actin structures and BAR domain proteins can regulate the tubular shape of spine necks, we implemented their net effect in our model by including spontaneous deviatoric curvature in the energy density of the system (Eq. 3) [78, 81–84].

Consider a tubular membrane with radius r and a spontaneous deviatoric curvature D_m_ along the neck with total length *l* (Fig. 1A), the equilibrium radius in the presence of spontaneous deviatoric curvature is given by 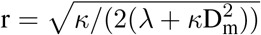 (Eq. S41). Since this radius depends on both the value of tension and the spontaneous deviatoric curvature (Fig. 5A), we define an effective tension 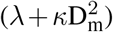. As a result, the relationship between neck radius, spontaneous deviatoric curvature, and tension in Fig. 5A collapses onto a single curve (Fig. S6B) as a function of this effective tension. Simulations confirm that the radii of tubular necks obtained from numerical solutions collapse onto a single curve as a function of effective tension (Fig. 5B).

**Figure 5:**
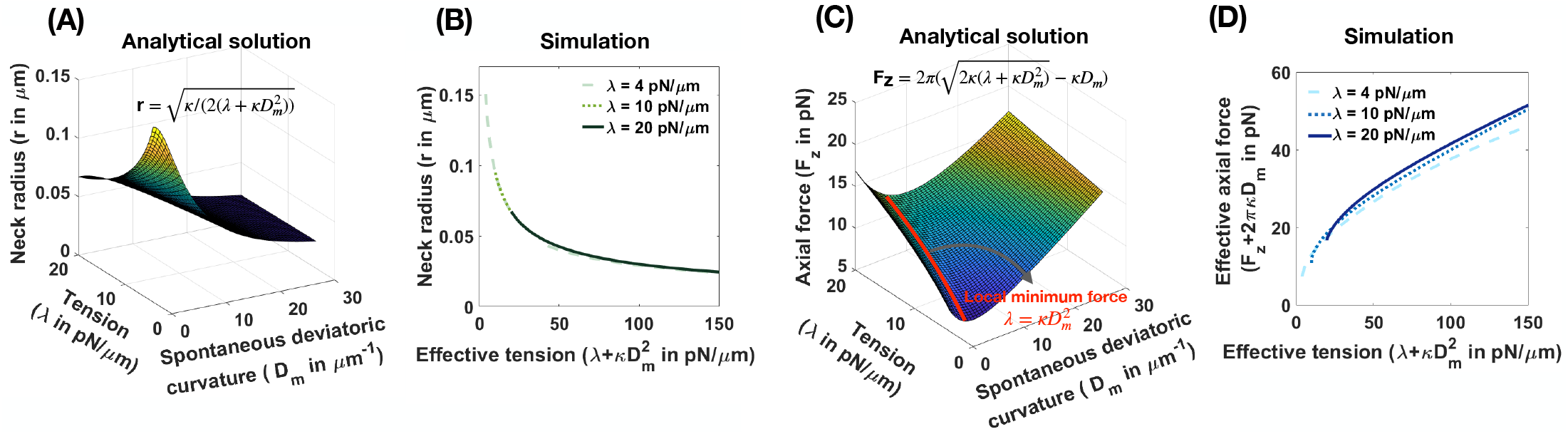
Effective tension including spontaneous deviatoric curvature regulates the neck radius and the magnitude of axial force in a tubular membrane. (A) Analytical solution for the neck radius of a tubular membrane as a function of spontaneous deviatoric curvature and tension 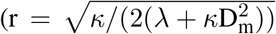, Eq. S41). (B) The neck radius obtained from numerical solutions as a function of effective tension 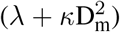. Here, for fixed three different tensions, we varied the effective tension by changing the spontaneous deviatoric curvature between 0 *<* D_m_ *<* 30 *µ*m^−1^. The radii of the membrane necks collapse onto a single curve for different tensions. (C) Analytical solution for the magnitude of an axial force needed to maintain a tubular protrusion as a function of spontaneous deviatoric curvature and tension 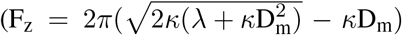, Eq. S41). The axial force needed to maintain a tubular protrusion has a local minimum along the red line where 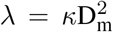 (Eq. S43). (D) The effective axial force (F_z_ + 2*πκ*D_m_) obtained from numerical solutions as a function of effective tension 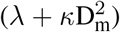. Here, for fixed three different tensions, we varied the effective tension by changing the spontaneous deviatoric curvature between 0 *<* D_m_ *<* 30 *µ*m^−1^. Effective axial forces collapse onto a single curve for different tensions.

Similarly, the axial force required to maintain a tubular membrane with radius r and spontaneous deviatoric curvature D_m_ along the total length L, is given by 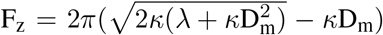 (Eq. S41). In Fig. 5C, we plotted the axial force as a function of tension and spontaneous deviatoric curvature. We found that the axial force has a local minimum along the red line (Fig. 5C) where 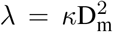 (Eq. S42) and F_z,min_ = 2*πκ*D_m_ (Eq. S42). The 3D surface in Fig. 5C can be reduced to a single curve by defining the effective axial force as F_z_ + 2*πκ*D_m_ and plotting it as a function of effective tension (Fig. S6D). We also plotted the effective axial force obtained from numerical solutions as a function of effective tension (Fig. 5D). We observed that consistent with the analytical prediction, for different tensions, the effective axial forces collapse onto a single curve as a function of effective tension (Fig. 5D). These results suggest that effective tension 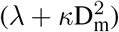 regulates the radius of dendritic spine necks.

### 3.5 Cooperation of forces and induced spontaneous deviatoric curvature offers multiple pathways for spine shape maintenance

Thus far, we have focused on the role of forces (axial and normal) on spine head shape and the role of spontaneous deviatoric curvature representing periodic rings on the spine neck radius. Next, we asked if the cooperation of these two different mechanisms could further influence the spine geometries and the energy landscape associated with these features. In other words, we asked if the combination of spontaneous deviatoric curvature and applied forces could result in lower energy states for the same spine geometry. To answer this question, we sought to identify the parameters that give rise to thin spines with the same geometric parameters. We explain this approach with a specific example below.

As noted before, when only normal forces are used, a normal force of F_n_ = 168.18 pN under a tension of *λ* = 36 pN*/µ*m is required to form a thin spine with a neck radius of r = 0.05 *µ*m and head volume of V = 0.033 *µ*m^3^ (Fig. 6A, left). We can also obtain a thin spine with the same dimensions, by using a prescribed spontaneous deviatoric curvature D_m_ = 10 *µ*m^−1^ along the neck and an applied force of F_n_ = 43 pN along the head for *λ* = 10 pN*/µ*m (Fig. 6A, right). Thus, for the same shape parameters, in the presence of spontaneous deviatoric curvature, the value of force required is roughly a quarter of the force required in the absence of spontaneous deviatoric curvature (Fig. 6A). Similarly, when a combination of axial force along the spine head and spontaneous deviatoric curvature along the neck is used, a thin spine with r ∼ 0.05 *µ*m and head volume V ∼ 0.033 *µ*m^3^ can be formed with F_z_ = 7.71 pN and spontaneous deviatoric curvature D_m_ = 10 *µ*m^−1^ when *λ* = 10 pN*/µ*m (Fig. 6B). Thus, in both these cases (axial and normal forces) for the formation of thin spines, we note that access to spontaneous deviatoric curvature significantly reduces the forces required to form and maintain thin spines.

**Figure 6:**
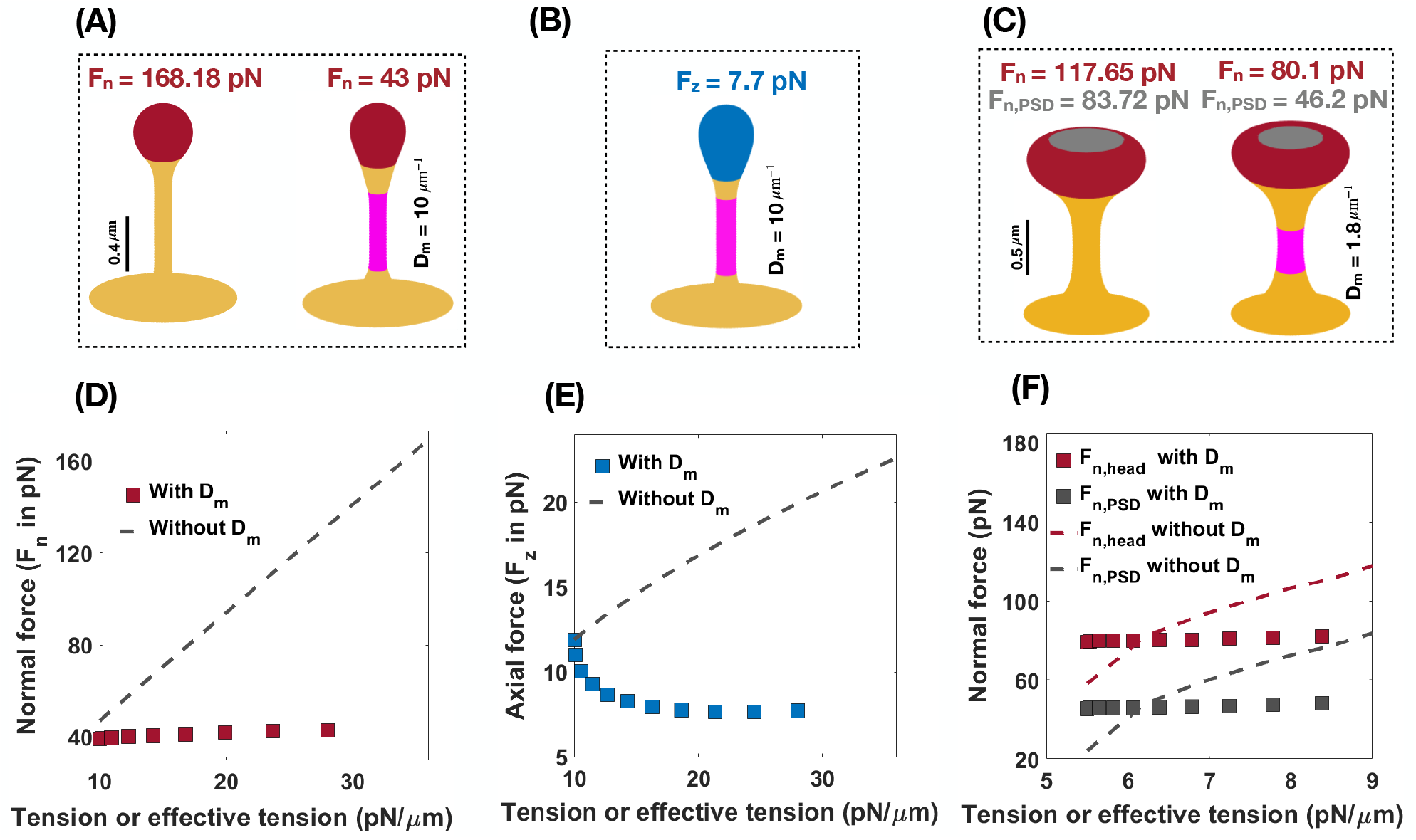
Formation of thin and mushroom shaped spines with a combination of forces and spontaneous deviatoric curvature. (A) Formation of a thin-shaped spine by applying a uniform normal force density along the spine head (left) versus applying a uniform normal force density along the head and spontaneous deviatoric curvature (purple region) along the spine neck (right) (*λ* =10 pN*/µ*m). (B) Formation of a thin-shaped spine by applying an axial force along the spherical cap (blue region) and spontaneous deviatoric curvature along the spine neck (purple region), *λ* =10 pN*/µ*m. All thin spines in panels A and B have a neck radius r ∼ 0.05 *µ*m and head volume V ∼ 0.033 *µ*m^3^. (C) Formation of a mushroom-shaped spine by applying a non-uniform normal force density along the spine head (left) versus applying a non-uniform normal force density along the head and spontaneous deviatoric curvature along the spine neck (purple region), (right), *λ* =5.5 pN*/µ*m. The formed mushroom spine with normal forces F_n_ = 80.1 pN and F_n,PSD_ = 46.2 pN and deviatoric curvature D_m_ = 1.8 *µ*m^−1^ has a neck radius r ∼ 0.1 *µ*m and head volume V ∼ 0.27 *µ*m^3^. (D) The magnitude of a normal force that is required to form a thin-shaped spine with and without spontaneous deviatoric curvature as a function of effective tension and tension, respectively. (E) The magnitude of an axial force that is required to form a thin-shaped spine with and without spontaneous deviatoric curvature as a function of effective tension and tension, respectively. (F) The magnitude of normal forces in the spine head and in PSD that is required to form a mushroom spine with and without spontaneous deviatoric curvature as a function of effective tension and tension, respectively.

Not surprisingly, these same results hold for mushroom-shaped spines too. As we have shown before, to form a mushroom spine with a neck radius of r = 0.1 *µ*m and head volume of V ∼ 0.25 *µ*m^3^, normal forces of F_n_ = 117.65 pN along the spine head and F_n,PSD_ = 83.72 pN along the PSD are required under a tension of *λ* = 9 pN*/µ*m (Fig. 6C, left). We can also form a mushroom spine with the same dimensions and lower tension (*λ* = 5.5 pN*/µ*m) by prescribing a spontaneous deviatoric curvature D_m_ = 1.8 *µ*m^−1^ along the spine neck and normal forces of F_n_ = 80.1 pN and F_n,PSD_ = 45.17 pN along the spine head and PSD, respectively (Fig. 6C, right).

In Figs. 6D-F, we plotted the magnitude of forces that are required to form thin and mushroom-shaped spines with or without spontaneous deviatoric curvature as a function of tension alone (with no spontaneous deviatoric curvature) or effective tension (with spontaneous deviatoric curvature). We observed that with increasing effective tension, the magnitude of the normal force that is required to form a thin spine with spontaneous deviatoric curvature (red squares) is almost constant (Fig. 6D). However, the magnitude of the normal force that is needed to form a thin spine without spontaneous deviatoric curvature (dashed line) increases linearly with increasing tension (Eq.6 and Fig. 6D). In the case of the formation of a thin spine with an axial force, we found that in the presence of spontaneous deviatoric curvature, the magnitude of axial force (blue squares) decreases slightly and then becomes constant with increasing effective tension (Fig. 6E). In contrast, without spontaneous deviatoric curvature, the magnitude of axial force (dashed line) increases with increasing tension (Fig. 6E). Similar to the thin-shaped spine, with spontaneous deviatoric curvature along the spine neck, the magnitude of normal forces in the head (red square) and PSD (gray square) region that are required to form a mushroom spine is almost constant with increasing effective tension (Fig. 6F). However, without spontaneous deviatoric curvature, the magnitude of forces in both regions increases with increasing tension (Fig. 6F).

To further compare thin and mushroom spines shown in Fig. 6, we computed the components of energy (Eq. 1) and the total energy of the system for each shape (Tables S2 & S3 and Figs. 7C & D). Based on our results, by prescribing spontaneous deviatoric curvature D_m_ along the spine neck, the bending energy due to deviatoric curvature decreases (Tables. S2 & S3). This is because the deviatoric curvature *D* along the neck tends to D_m_ and minimizes the bending energy (Tables S2 & S3). Additionally, in the presence of spontaneous deviatoric curvature, in our simulation, we set the tension to lower values compared to the condition that D_m_ = 0. Therefore, the work that is done by tension and forces to bend the membrane reduces for the case that the spines obtained with a combination of force and spontaneous deviatoric curvature (Tables. S2 & S3). For example, to form a thin spine shown in Fig. 6, the work that is done by an axial force with a spontaneous deviatoric curvature (Fig. 6B) is almost one third of the work that is done by a normal force without spontaneous deviatoric curvature (Fig. 6A and Tables. S2 & S3).

**Figure 7:**
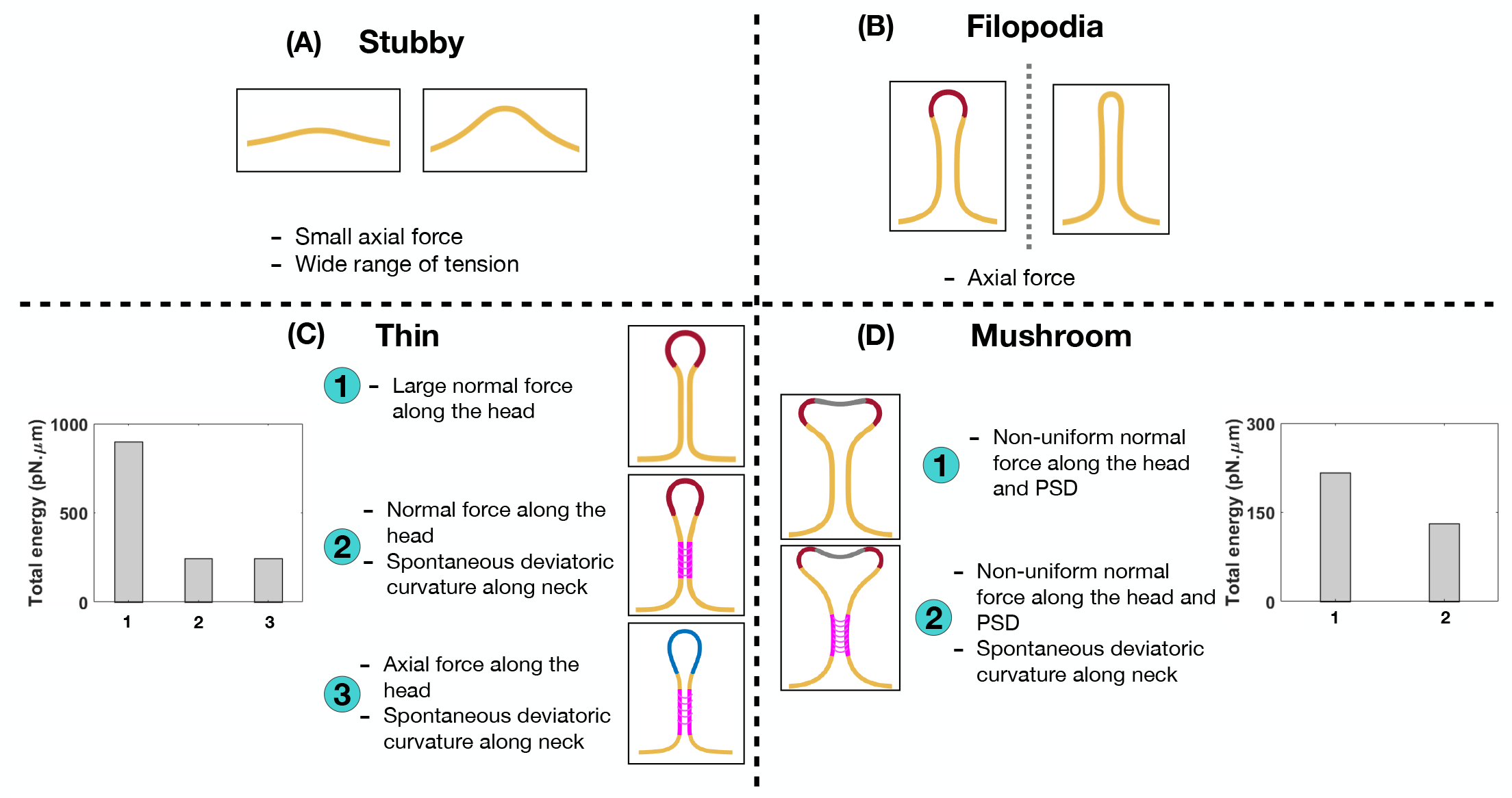
Characterizing different shapes of dendritic spines based on the mechanical model. (A) Stubby spines can be formed with an axial force and in a wide range of tensions. (B) An axial force is sufficient to form a long filopodial spine. (C) A thin-shaped spine can be formed with three different mechanisms; (1) a uniform normal force density along the spine head, (2) a uniform normal force density along the spine head and spontaneous deviatoric curvature along the neck, and (3) a uniform axial force density along the spine head and spontaneous deviatoric curvature along the neck. In the bar plot, the total energy of the system is shown for three different mechanisms. The total energy of the system for the second and third mechanisms with spontaneous deviatoric curvature is much less than the energy for the first mechanism with just a normal force. (D) A mushroom-shaped spine can be formed with two different mechanisms; (1) a non-uniform normal force density along the spine head and PSD region and (2) a non-uniform normal force density along the spine head and PSD region plus a spontaneous deviatoric curvature along the spine neck. The resulting mushroom spine with a combination of normal forces and spontaneous deviatoric curvature has lower energy compared to the spine that is formed with just normal forces (bar graph).

In the bar plots of Figs. 7C &D, we compared the total energy of thin and mushroom spines formed with different mechanisms. We observed that in both thin and mushroom spines, the total energy of the system dramatically decreases when the spines form with a combination of forces and spontaneous deviatoric curvature (Figs. 7C &D). This result suggests that spontaneous deviatoric curvature can alter the energy landscape of thin and mushroom dendritic spines to a lower energy state.

## 4 Discussion

Previously, we showed that the coupled dynamics of signaling and actin remodeling can alter spine volume in a biophysical model [67] without considering the geometry of the spines or considering the role of spine shape in regulating different signaling pathways [114–118]. In this work, we present a simplified mechanical model for studying the role of different force distributions and energy contributions that are associated with the different spine shapes noted in the literature. Our results show that different spine shapes can be associated with different forces and spontaneous deviatoric curvature distributions, giving us insight into the mechanical design principles of spine formation and maintenance (Fig. 7).

We show that stubby spines can be formed for a wide range of tensions and low forces (Fig. 2). From a spine formation viewpoint, this makes sense, since during development the stubby spines can be the initial protrusions that form out of the dendrites. Given the ubiquitous nature of stubby spines [4, 12, 19], our results suggest that the prevalence of stubby spines could be due to the mechanical ease which they can be formed. They may also represent a temporarily stable state adopted by shrinking spines during synapse removal. Filopodia have the same force-length and force-radius relationships as membrane tubules that can be formed with micropipettes [119], optical tweezers [120], or by kinesin motor proteins [121] (Fig. 2). Based on our results, dendritic filopodia can be formed with a relatively small axial force, which make them good candidates as initial protrusions for the formation of mature thin and mushroom spines. Thin and mushroom spines, which have defined head shapes, require more mechanical features – heterogeneous force distributions, normal or axial forces, and an induced spontaneous deviatoric curvature representing the periodic protein rings or other deviatoric curvature inducing mechanics along the neck.

In the case of thin spines, we find that the mechanical design principles that support the formation of a spherical head are (1) large normal force along the head (Fig. 3), (2) normal force along the head with a spontaneous deviatoric curvature along the neck (Fig. 6A), and (3) an axial force along the head with a spontaneous deviatoric curvature along the neck (Fig. 6B). Within these mechanisms, the presence of spontaneous deviatoric curvature significantly reduces the total energy of the spine (Fig. 7C). Similarly, for mushroom spines, in addition to non-uniform forces along the head and the PSD (Fig. 4), the spine can be formed with a combination of forces in the head and spontaneous deviatoric curvature along the neck (Fig. 6C) while the spontaneous deviatoric curvature results in a lower energy state (Fig. 7D).

These findings have implications for our understanding of how mechanical aspects of membrane dynamics such as bending, tension, membrane-protein interactions, and interactions of the membrane with the cytoskeleton play critical roles in spine geometry maintenance, particularly in structural plasticity. Many of the events associated with synaptic plasticity alter spine size and shape through changes in F-actin dynamics and the dynamics of the actin related proteins [38–40]. The net impact of changes in actin remodeling would likely result in changes in force distribution. Another important and, as yet, under explored aspect of synaptic plasticity is the role of cortical membrane tension, including the effect of the membrane in-plane stresses and membrane-cytoskeleton interactions. We know that spines are sites of active vesicle trafficking events, such as endo- and exocytosis, and that these processes alter the membrane surface area and thereby alter the membrane tension [122, 123]. Here, we show that the effective membrane tension can play an important role in altering the energy required for the maintenance of different spine shapes. One of the main impacts of such effective tension is that because of the cooperative effects of spontaneous deviatoric curvature and the applied forces, the energy required to maintain certain spine shapes may be lower. Thus, we show that there are different mechanical pathways that are likely associated with the different spine shapes and that some mechanisms may be energetically more favorable than others.

Despite these insights, our model has certain limitations. We do not explicitly consider the remodeling of the actin network or the dynamics of the associated proteins, but use force as a lumped parameter. Additionally, the use of axisymmetric coordinates restricts our ability to obtain realistic spine shapes [124]. We note that the technology required to simulate these processes is quite challenging [125] and is ongoing research in our group.

The impact of mechanical aspects of actin remodeling and membrane mechanics on structural plasticity is highly intriguing and we are only beginning to understand their effects on spine functionality. This complexity is immediately apparent in dendritic spines, which undergo dynamic changes, both mechanical and biochemical during structural plasticity spatiotemporal scales. Our minimal model provides insights into the possible mechanical aspects underlying the characteristic geometries associated with dendritic spines. This is an important step towards deciphering the intricate mechanochemistry of structural plasticity and dendritic spine development.

## Supporting information

Supplementary material for "Mechanical principles governing dendritic spine shapes"

## Acknowledgments

We would like to thank Dr. Christopher Lee, Can Uysalel, Arijit Mahapatra, and Andrew Nguyen for critical discussions and feedback on the manuscript. This work was supported by Air Force Office of Scientific Research FA9550-18-1-0051 to P.R and M.K.B. was supported by a National Defense Science and Engineering Graduate (NDSEG) Fellowship.

## Author contributions

H.A. and P.R. conceived the research, H.A., M.K.B, and P.R. conducted the research and analyzed the data, H.A., M.K.B, S.H. and P.R. wrote the paper. All authors reviewed the manuscript and agreed on the contents of the paper.

## References

[1] J. Nishiyama, “Plasticity of dendritic spines: Molecular function and dysfunction in neurodevelopmental disorders,” Psychiatry and Clinical Neurosciences, 2019.

[2] Y. Nakahata and R. Yasuda, “Plasticity of spine structure: local signaling, translation and cytoskeletal reorganization,” Frontiers in synaptic neuroscience, vol. 10, p. 29, 2018.

[3] M. Bosch and Y. Hayashi, “Structural plasticity of dendritic spines,” Current opinion in neurobiology, vol. 22, no. 3, pp. 383–388, 2012.

[4] E. G. Gray, “Electron microscopy of synaptic contacts on dendrite spines of the cerebral cortex,” Nature, vol. 183, no. 4675, pp. 1592–1593, 1959.

[5] G. M. Shepherd, “The dendritic spine: a multifunctional integrative unit,” Journal of neurophysiology, vol. 75, no. 6, pp. 2197–2210, 1996.

[6] K. M. Harris and S. Kater, “Dendritic spines: cellular specializations imparting both stability and flexibility to synaptic function,” Annual review of neuroscience, vol. 17, no. 1, pp. 341–371, 1994.

[7] J. I. Arellano, R. Benavides-Piccione, J. DeFelipe, and R. Yuste, “Ultrastructure of dendritic spines: correlation between synaptic and spine morphologies,” Frontiers in neuroscience, vol. 1, p. 10, 2007.

[8] M. Patterson and R. Yasuda, “Signalling pathways underlying structural plasticity of dendritic spines,” British journal of pharmacology, vol. 163, no. 8, pp. 1626–1638, 2011.

[9] F. Engert and T. Bonhoeffer, “Dendritic spine changes associated with hippocampal long-term synaptic plasticity,” Nature, vol. 399, no. 6731, p. 66, 1999.

[10] M. F. Bear and R. C. Malenka, “Synaptic plasticity: Ltp and ltd,” Current opinion in neurobiology, vol. 4, no. 3, pp. 389–399, 1994.

[11] K. M. Harris, J. C. Fiala, and L. Ostroff, “Structural changes at dendritic spine synapses during long-term potentiation,” Philosophical Transactions of the Royal Society of London. Series B: Biological Sciences, vol. 358, no. 1432, pp. 745–748, 2003.

[12] K. M. Harris, “Structure, development, and plasticity of dendritic spines,” Current opinion in neurobiology, vol. 9, no. 3, pp. 343–348, 1999.

[13] L. E. Ostroff, J. C. Fiala, B. Allwardt, and K. M. Harris, “Polyribosomes redistribute from dendritic shafts into spines with enlarged synapses during ltp in developing rat hippocampal slices,” Neuron, vol. 35, no. 3, pp. 535–545, 2002.

[14] L. J. Petrak, K. M. Harris, and S. A. Kirov, “Synaptogenesis on mature hippocampal dendrites occurs via filopodia and immature spines during blocked synaptic transmission,” Journal of Comparative Neurology, vol. 484, no. 2, pp. 183–190, 2005.

[15] E. Fifkova, “A possible mechanism of morphometric changes in dendritic spines induced by stimulation,” Cellular and molecular neurobiology, vol. 5, no. 1-2, pp. 47–63, 1985.

[16] B. Calabrese, M. S. Wilson, and S. Halpain, “Development and regulation of dendritic spine synapses,” Physiology, vol. 21, no. 1, pp. 38–47, 2006.

[17] A. Peters and I. R. Kaiserman-Abramof, “The small pyramidal neuron of the rat cerebral cortex. the perikaryon, dendrites and spines,” American Journal of Anatomy, vol. 127, no. 4, pp. 321–355, 1970.

[18] K. M. Harris, F. E. Jensen, and B. Tsao, “Three-dimensional structure of dendritic spines and synapses in rat hippocampus (ca1) at postnatal day 15 and adult ages: implications for the maturation of synaptic physiology and long-term potentiation [published erratum appears in j neurosci 1992 aug; 12 (8): following table of contents],” Journal of Neuroscience, vol. 12, no. 7, pp. 2685–2705, 1992.

[19] J. C. Fiala, M. Feinberg, V. Popov, and K. M. Harris, “Synaptogenesis via dendritic filopodia in developing hippocampal area ca1,” Journal of Neuroscience, vol. 18, no. 21, pp. 8900–8911, 1998.

[20] A. J. Holtmaat, J. T. Trachtenberg, L. Wilbrecht, G. M. Shepherd, X. Zhang, G. W. Knott, and K. Svoboda, “Transient and persistent dendritic spines in the neocortex in vivo,” Neuron, vol. 45, no. 2, pp. 279–291, 2005.

[21] Y. Zuo, A. Lin, P. Chang, and W.-B. Gan, “Development of long-term dendritic spine stability in diverse regions of cerebral cortex,” Neuron, vol. 46, no. 2, pp. 181–189, 2005.

[22] H. Kasai, M. Matsuzaki, J. Noguchi, N. Yasumatsu, and H. Nakahara, “Structure–stability–function relationships of dendritic spines,” Trends in neurosciences, vol. 26, no. 7, pp. 360–368, 2003.

[23] J. Bourne and K. M. Harris, “Do thin spines learn to be mushroom spines that remember?,” Current opinion in neurobiology, vol. 17, no. 3, pp. 381–386, 2007.

[24] K. P. Berry and E. Nedivi, “Spine dynamics: are they all the same?,” Neuron, vol. 96, no. 1, pp. 43–55, 2017.

[25] J. Grutzendler, N. Kasthuri, and W.-B. Gan, “Long-term dendritic spine stability in the adult cortex,” Nature, vol. 420, no. 6917, pp. 812–816, 2002.

[26] J. T. Trachtenberg, B. E. Chen, G. W. Knott, G. Feng, J. R. Sanes, E. Welker, and K. Svoboda, “Long-term in vivo imaging of experience-dependent synaptic plasticity in adult cortex,” Nature, vol. 420, no. 6917, pp. 788–794, 2002.

[27] M. Matsuzaki, G. C. Ellis-Davies, T. Nemoto, Y. Miyashita, M. Iino, and H. Kasai, “Dendritic spine geometry is critical for ampa receptor expression in hippocampal ca1 pyramidal neurons,” Nature neuroscience, vol. 4, no. 11, p. 1086, 2001.

[28] O. Ganeshina, R. Berry, R. Petralia, D. Nicholson, and Y. Geinisman, “Synapses with a segmented, completely partitioned postsynaptic density express more ampa receptors than other axospinous synaptic junctions,” Neuroscience, vol. 125, no. 3, pp. 615–623, 2004.

[29] M. C. Ashby, S. R. Maier, A. Nishimune, and J. M. Henley, “Lateral diffusion drives constitutive exchange of ampa receptors at dendritic spines and is regulated by spine morphology,” Journal of Neuroscience, vol. 26, no. 26, pp. 7046–7055, 2006.

[30] R. Kanjhan, P. G. Noakes, and M. C. Bellingham, “Emerging roles of filopodia and dendritic spines in motoneuron plasticity during development and disease,” Neural plasticity, vol. 2016, 2016.

[31] R. Yuste and T. Bonhoeffer, “Genesis of dendritic spines: insights from ultrastructural and imaging studies,” Nature Reviews Neuroscience, vol. 5, no. 1, p. 24, 2004.

[32] A. Rodriguez, D. B. Ehlenberger, D. L. Dickstein, P. R. Hof, and S. L. Wearne, “Automated three-dimensional detection and shape classification of dendritic spines from fluorescence microscopy images,” PloS one, vol. 3, no. 4, p. e1997, 2008.

[33] J. Spacek and K. M. Harris, “Three-dimensional organization of smooth endoplasmic reticulum in hippocampal ca1 dendrites and dendritic spines of the immature and mature rat,” Journal of Neuroscience, vol. 17, no. 1, pp. 190–203, 1997.

[34] M. E. Dailey and S. J. Smith, “The dynamics of dendritic structure in developing hippocampal slices,” Journal of neuroscience, vol. 16, no. 9, pp. 2983–2994, 1996.

[35] M. Miller and A. Peters, “Maturation of rat visual cortex. ii. a combined golgi-electron microscope study of pyramidal neurons,” Journal of Comparative Neurology, vol. 203, no. 4, pp. 555–573, 1981.

[36] N. E. Ziv and S. J. Smith, “Evidence for a role of dendritic filopodia in synaptogenesis and spine formation,” Neuron, vol. 17, no. 1, pp. 91–102, 1996.

[37] A. J. Koleske, “Molecular mechanisms of dendrite stability,” Nature Reviews Neuroscience, vol. 14, no. 8, pp. 536–550, 2013.

[38] E. Bertling and P. Hotulainen, “New waves in dendritic spine actin cytoskeleton: From branches and bundles to rings, from actin binding proteins to post-translational modifications,” Molecular and Cellular Neuroscience, vol. 84, pp. 77–84, 2017.

[39] P. Hotulainen and C. C. Hoogenraad, “Actin in dendritic spines: connecting dynamics to function,” The Journal of cell biology, vol. 189, no. 4, pp. 619–629, 2010.

[40] D. Landis and T. S. Reese, “Cytoplasmic organization in cerebellar dendritic spines.,” The Journal of cell biology, vol. 97, no. 4, pp. 1169–1178, 1983.

[41] C. S. Peskin, G. M. Odell, and G. F. Oster, “Cellular motions and thermal fluctuations: the brownian ratchet,” Biophysical journal, vol. 65, no. 1, pp. 316–324, 1993.

[42] A. Mogilner and G. Oster, “Cell motility driven by actin polymerization,” Biophysical journal, vol. 71, no. 6, pp. 3030–3045, 1996.

[43] C. Miermans, R. Kusters, C. Hoogenraad, and C. Storm, “Biophysical model of the role of actin remodeling on dendritic spine morphology,” PloS one, vol. 12, no. 2, p. e0170113, 2017.

[44] N. A. Frost, H. Shroff, H. Kong, E. Betzig, and T. A. Blanpied, “Single-molecule discrimination of discrete perisynaptic and distributed sites of actin filament assembly within dendritic spines,” Neuron, vol. 67, no. 1, pp. 86–99, 2010.

[45] F. Korobova and T. Svitkina, “Molecular architecture of synaptic actin cytoskeleton in hippocampal neurons reveals a mechanism of dendritic spine morphogenesis,” Molecular biology of the cell, vol. 21, no. 1, pp. 165–176, 2010.

[46] S. Nanguneri, R. Pramod, N. Efimova, D. Das, M. Jose, T. Svitkina, and D. Nair, “Characterization of nanoscale organization of f-actin in morphologically distinct dendritic spines in vitro using supervised learning,” eNeuro, vol. 6, no. 4, 2019.

[47] S. Halpain, “Actin and the agile spine: how and why do dendritic spines dance?,” Trends in neurosciences, vol. 23, no. 4, pp. 141–146, 2000.

[48] A. Rao and A. M. Craig, “Signaling between the actin cytoskeleton and the postsynaptic density of dendritic spines,” Hippocampus, vol. 10, no. 5, pp. 527–541, 2000.

[49] T. Tada and M. Sheng, “Molecular mechanisms of dendritic spine morphogenesis,” Current opinion in neurobiology, vol. 16, no. 1, pp. 95–101, 2006.

[50] P. Hotulainen, O. Llano, S. Smirnov, K. Tanhuanpää, J. Faix, C. Rivera, and P. Lappalainen, “Defining mechanisms of actin polymerization and depolymerization during dendritic spine morphogenesis,” The Journal of cell biology, vol. 185, no. 2, pp. 323–339, 2009.

[51] M. Bucher, T. Fanutza, and M. Mikhaylova, “Cytoskeletal makeup of the synapse: Shaft versus spine,” Cytoskeleton.

[52] J. Bär, O. Kobler, B. Van Bommel, and M. Mikhaylova, “Periodic f-actin structures shape the neck of dendritic spines,” Scientific reports, vol. 6, p. 37136, 2016.

[53] á. Racz and R. Weinberg, “Spatial organization of cofilin in dendritic spines,” Neuroscience, vol. 138, no. 2, pp. 447–456, 2006.

[54] B. Rácz and R. J. Weinberg, “Microdomains in forebrain spines: an ultrastructural perspective,” Molecular neurobiology, vol. 47, no. 1, pp. 77–89, 2013.

[55] W.-H. Lin and D. J. Webb, “Actin and actin-binding proteins: masters of dendritic spine formation, morphology, and function,” The open neuroscience journal, vol. 3, p. 54, 2009.

[56] N. Honkura, M. Matsuzaki, J. Noguchi, G. C. Ellis-Davies, and H. Kasai, “The subspine organization of actin fibers regulates the structure and plasticity of dendritic spines,” Neuron, vol. 57, no. 5, pp. 719–729, 2008.

[57] A. R. Huehn, J. P. Bibeau, A. C. Schramm, W. Cao, M. Enrique, and C. V. Sindelar, “Structures of cofilin-induced structural changes reveal local and asymmetric perturbations of actin filaments,” Proceedings of the National Academy of Sciences, 2020.

[58] A. Chazeau and G. Giannone, “Organization and dynamics of the actin cytoskeleton during dendritic spine morphological remodeling,” Cellular and molecular life sciences, vol. 73, no. 16, pp. 3053–3073, 2016.

[59] P. J. Goldschmidt-Clermont, L. M. Machesky, S. K. Doberstein, and T. D. Pollard, “Mechanism of the interaction of human platelet profilin with actin.,” The Journal of cell biology, vol. 113, no. 5, pp. 1081–1089, 1991.

[60] L. A. Helgeson and B. J. Nolen, “Mechanism of synergistic activation of arp2/3 complex by cortactin and n-wasp,” Elife, vol. 2, p. e00884, 2013.

[61] N. Efimova, F. Korobova, M. C. Stankewich, A. H. Moberly, D. B. Stolz, J. Wang, A. Kashina, M. Ma, and T. Svitkina, “*β*iii spectrin is necessary for formation of the constricted neck of dendritic spines and regulation of synaptic activity in neurons,” Journal of Neuroscience, vol. 37, no. 27, pp. 6442–6459, 2017.

[62] M. A. Mikati, E. E. Grintsevich, and E. Reisler, “Drebrin-induced stabilization of actin filaments,” Journal of Biological Chemistry, vol. 288, no. 27, pp. 19926–19938, 2013.

[63] M. Kneussel and W. Wagner, “Myosin motors at neuronal synapses: drivers of membrane transport and actin dynamics,” Nature Reviews Neuroscience, vol. 14, no. 4, pp. 233–247, 2013.

[64] K.-I. Okamoto, R. Narayanan, S. H. Lee, K. Murata, and Y. Hayashi, “The role of camkii as an f-actin-bundling protein crucial for maintenance of dendritic spine structure,” Proceedings of the National Academy of Sciences, vol. 104, no. 15, pp. 6418–6423, 2007.

[65] J. Borovac, M. Bosch, and K. Okamoto, “Regulation of actin dynamics during structural plasticity of dendritic spines: Signaling messengers and actin-binding proteins,” Molecular and Cellular Neuroscience, 2018.

[66] S. Khan, Y. Zou, A. Amjad, A. Gardezi, C. L. Smith, C. Winters, and T. S. Reese, “Sequestration of camkii in dendritic spines in silico,” Journal of computational neuroscience, vol. 31, no. 3, pp. 581–594, 2011.

[67] P. Rangamani, M. G. Levy, S. Khan, and G. Oster, “Paradoxical signaling regulates structural plasticity in dendritic spines,” Proceedings of the National Academy of Sciences, vol. 113, no. 36, pp. E5298–E5307, 2016.

[68] B. Calabrese and S. Halpain, “Essential role for the pkc target marcks in maintaining dendritic spine morphology,” Neuron, vol. 48, no. 1, pp. 77–90, 2005.

[69] I. Hlushchenko, M. Koskinen, and P. Hotulainen, “Dendritic spine actin dynamics in neuronal maturation and synaptic plasticity,” Cytoskeleton, vol. 73, no. 9, pp. 435–441, 2016.

[70] J. Saarikangas, N. Kourdougli, Y. Senju, G. Chazal, M. Segerstråle, R. Minkeviciene, J. Kuurne, P. K. Mattila, L. Garrett, S. M. Hölter, et al., “Mim-induced membrane bending promotes dendritic spine initiation,” Developmental cell, vol. 33, no. 6, pp. 644–659, 2015.

[71] B. R. Carlson, K. E. Lloyd, A. Kruszewski, I.-H. Kim, R. M. Rodriguiz, C. Heindel, M. Faytell, S. M. Dudek, W. C. Wetsel, and S. H. Soderling, “Wrp/srgap3 facilitates the initiation of spine development by an inverse f-bar domain, and its loss impairs long-term memory,” Journal of Neuroscience, vol. 31, no. 7, pp. 2447–2460, 2011.

[72] M. M. Kessels and B. Qualmann, “Different functional modes of bar domain proteins in formation and plasticity of mammalian postsynapses,” J Cell Sci, vol. 128, no. 17, pp. 3177–3185, 2015.

[73] Y. Wakita, T. Kakimoto, H. Katoh, and M. Negishi, “The f-bar protein rapostlin regulates dendritic spine formation in hippocampal neurons,” Journal of Biological Chemistry, vol. 286, no. 37, pp. 32672–32683, 2011.

[74] B. J. Peter, H. M. Kent, I. G. Mills, Y. Vallis, P. J. G. Butler, P. R. Evans, and H. T. McMahon, “Bar domains as sensors of membrane curvature: the amphiphysin bar structure,” Science, vol. 303, no. 5657, pp. 495–499, 2004.

[75] A. Frost, V. M. Unger, and P. De Camilli, “The bar domain superfamily: membrane-molding macromolecules,” Cell, vol. 137, no. 2, pp. 191–196, 2009.

[76] A. Frost, R. Perera, A. Roux, K. Spasov, O. Destaing, E. H. Egelman, P. De Camilli, and V. M. Unger, “Structural basis of membrane invagination by f-bar domains,” Cell, vol. 132, no. 5, pp. 807–817, 2008.

[77] A. Shimada, H. Niwa, K. Tsujita, S. Suetsugu, K. Nitta, K. Hanawa-Suetsugu, R. Akasaka, Y. Nishino, M. Toyama, L. Chen, et al., “Curved efc/f-bar-domain dimers are joined end to end into a filament for membrane invagination in endocytosis,” Cell, vol. 129, no. 4, pp. 761–772, 2007.

[78] A. Iglič, B. Babnik, U. Gimsa, and V. Kralj-Iglič, “On the role of membrane anisotropy in the beading transition of undulated tubular membrane structures,” Journal of Physics A: Mathematical and General, vol. 38, no. 40, p. 8527, 2005.

[79] V. Kralj-Iglič, A. Iglič, H. Hägerstrand, and P. Peterlin, “Stable tubular microexovesicles of the erythrocyte membrane induced by dimeric amphiphiles,” Physical Review E, vol. 61, no. 4, p. 4230, 2000.

[80] A. Iglič, B. Babnik, K. Bohinc, M. Fošnarič, H. Hägerstrand, and V. Kralj-Iglič, “On the role of anisotropy of membrane constituents in formation of a membrane neck during budding of a multicomponent membrane,” Journal of biomechanics, vol. 40, no. 3, pp. 579–585, 2007.

[81] V. Kralj-Iglič, V. Heinrich, S. Svetina, and B. Ž ekš, “Free energy of closed membrane with anisotropic inclusions,” The European Physical Journal B-Condensed Matter and Complex Systems, vol. 10, no. 1, pp. 5–8, 1999.

[82] V. Kralj-Iglič, S. Svetina, and B. Ž ekž, “Shapes of bilayer vesicles with membrane embedded molecules,” European biophysics journal, vol. 24, no. 5, pp. 311–321, 1996.

[83] A. Iglič, V. Kralj-Iglič, and J. Majhenc, “Cylindrical shapes of closed lipid bilayer structures correspond to an extreme area difference between the two monolayers of the bilayer,” Journal of biomechanics, vol. 32, no. 12, pp. 1343–1347, 1999.

[84] D. Kabaso, N. Bobrovska, W. Góźdź, N. Gov, V. Kralj-Iglič, P. Veranič, and A. Iglič, “On the role of membrane anisotropy and bar proteins in the stability of tubular membrane structures,” Journal of biomechanics, vol. 45, no. 2, pp. 231–238, 2012.

[85] M. Bonilla-Quintana, F. Wörgötter, C. Tetzlaff, and M. Fauth, “Modeling the shape of synaptic spines by their actin dynamics,” bioRxiv, p. 817932, 2019.

[86] H. Deuling and W. Helfrich, “Red blood cell shapes as explained on the basis of curvature elasticity,” Biophysical journal, vol. 16, no. 8, pp. 861–868, 1976.

[87] W. Helfrich, “Elastic properties of lipid bilayers: theory and possible experiments,” Zeitschrift fur Naturforschung C, vol. 28, no. 11-12, pp. 693–703, 1973.

[88] P. B. Canham, “The minimum energy of bending as a possible explanation of the biconcave shape of the human red blood cell,” Journal of theoretical biology, vol. 26, no. 1, pp. 61–81, 1970.

[89] A. Iglič, H. Hägerstrand, P. Veranič, A. Plemenitaš, and V. Kralj-Iglič, “Curvature-induced accumulation of anisotropic membrane components and raft formation in cylindrical membrane protrusions,” Journal of Theoretical Biology, vol. 240, no. 3, pp. 368–373, 2006.

[90] H. Alimohamadi and P. Rangamani, “Modeling membrane curvature generation due to membrane–protein interactions,” Biomolecules, vol. 8, no. 4, p. 120, 2018.

[91] W. Rawicz, K. Olbrich, T. McIntosh, D. Needham, and E. Evans, “Effect of chain length and unsaturation on elasticity of lipid bilayers,” Biophysical journal, vol. 79, no. 1, pp. 328–339, 2000.

[92] D. Steigmann, “Fluid films with curvature elasticity,” Archive for Rational Mechanics and Analysis, vol. 150, no. 2, pp. 127–152, 1999.

[93] P. Rangamani, A. Agrawal, K. K. Mandadapu, G. Oster, and D. J. Steigmann, “Interaction between surface shape and intra-surface viscous flow on lipid membranes,” Biomechanics and modeling in mechanobiology, pp. 1–13, 2013.

[94] H. Alimohamadi, B. Ovryn, and P. Rangamani, “Modeling membrane nanotube morphology: the role of heterogeneity in composition and material properties,” Scientific Reports, vol. 10, no. 1, pp. 1–15, 2020.

[95] A. Diz-Muñoz, D. A. Fletcher, and O. D. Weiner, “Use the force: membrane tension as an organizer of cell shape and motility,” Trends in cell biology, vol. 23, no. 2, pp. 47–53, 2013.

[96] H. Alimohamadi, A. S. Smith, R. B. Nowak, V. M. Fowler, and P. Rangamani, “Non-uniform distribution of myosin-mediated forces governs red blood cell membrane curvature through tension modulation,” PLOS Computational Biology, vol. 16, no. 5, p. e1007890, 2020.

[97] J. Weichsel and P. L. Geissler, “The more the tubular: Dynamic bundling of actin filaments for membrane tube formation,” PLoS computational biology, vol. 12, no. 7, p. e1004982, 2016.

[98] N. Walani, J. Torres, and A. Agrawal, “Endocytic proteins drive vesicle growth via instability in high membrane tension environment,” Proceedings of the National Academy of Sciences, vol. 112, no. 12, pp. E1423–E1432, 2015.

[99] E. Atilgan, D. Wirtz, and S. X. Sun, “Mechanics and dynamics of actin-driven thin membrane protrusions,” Biophysical journal, vol. 90, no. 1, pp. 65–76, 2006.

[100] J. E. Hassinger, G. Oster, D. G. Drubin, and P. Rangamani, “Design principles for robust vesiculation in clathrin-mediated endocytosis,” Proceedings of the National Academy of Sciences, vol. 114, no. 7, pp. E1118–E1127, 2017.

[101] A. Agrawal and D. J. Steigmann, “Modeling protein-mediated morphology in biomembranes,” Biomechanics and modeling in mechanobiology, vol. 8, no. 5, pp. 371–379, 2009.

[102] P. Rangamani, K. K. Mandadap, and G. Oster, “Protein-induced membrane curvature alters local membrane tension,” Biophysical journal, vol. 107, no. 3, pp. 751–762, 2014.

[103] N. Walani, J. Torres, and A. Agrawal, “Anisotropic spontaneous curvatures in lipid membranes,” Physical Review E, vol. 89, no. 6, p. 062715, 2014.

[104] M. Lokar, D. Kabaso, N. Resnik, K. Sepčić, V. Kralj-Iglič, P. Veranič, R. Zorec, and A. Iglič, “The role of cholesterol-sphingomyelin membrane nanodomains in the stability of intercellular membrane nanotubes,” International journal of nanomedicine, vol. 7, p. 1891, 2012.

[105] H. Alimohamadi, R. Vasan, J. Hassinger, J. C. Stachowiak, and P. Rangamani, “The role of traction in membrane curvature generation,” Mol. Biol. Cell, vol. 29, no. 16, pp. 2024–2035, 2018.

[106] B. Pontes, Y. Ayala, A. C. C. Fonseca, L. F. Romao, R. F. Amaral, L. T. Salgado, F. R. Lima, M. Farina, N. B. Viana, V. Moura-Neto, et al., “Membrane elastic properties and cell function,” PLoS One, vol. 8, no. 7, p. e67708, 2013.

[107] M. Bonilla-Quintana, F. Wörgötter, C. Tetzlaff, and M. Fauth, “Modeling the shape of synaptic spines by their actin dynamics,” Frontiers in Synaptic Neuroscience, vol. 12, p. 9, 2020.

[108] I. Derényi, F. Jülicher, and J. Prost, “Formation and interaction of membrane tubes,” Physical review letters, vol. 88, no. 23, p. 238101, 2002.

[109] T. R. Powers, G. Huber, and R. E. Goldstein, “Fluid-membrane tethers: minimal surfaces and elastic boundary layers,” Physical Review E, vol. 65, no. 4, p. 041901, 2002.

[110] G. G. Borisy and T. M. Svitkina, “Actin machinery: pushing the envelope,” Current opinion in cell biology, vol. 12, no. 1, pp. 104–112, 2000.

[111] M. Saleem, S. Morlot, A. Hohendahl, J. Manzi, M. Lenz, and A. Roux, “A balance between membrane elasticity and polymerization energy sets the shape of spherical clathrin coats,” Nature communications, vol. 6, no. 1, pp. 1–10, 2015.

[112] E. A. Evans, Mechanics and thermodynamics of biomembranes. CRC press, 2018.

[113] S. Ebrahimi and S. Okabe, “Structural dynamics of dendritic spines: molecular composition, geometry and functional regulation,” Biochimica et Biophysica Acta (BBA)-Biomembranes, vol. 1838, no. 10, pp. 2391–2398, 2014.

[114] M. Bell, T. Bartol, T. Sejnowski, and P. Rangamani, “Dendritic spine geometry and spine apparatus organization govern the spatiotemporal dynamics of calcium,” Journal of General Physiology, vol. 151, no. 8, pp. 1017–1034, 2019.

[115] D. Ohadi, D. L. Schmitt, B. Calabrese, S. Halpain, J. Zhang, and P. Rangamani, “Computational modeling reveals frequency modulation of calcium-camp/pka pathway in dendritic spines,” Biophysical Journal, vol. 117, no. 10, pp. 1963–1980, 2019.

[116] A. Cugno, T. M. Bartol, T. J. Sejnowski, R. Iyengar, and P. Rangamani, “Geometric principles of second messenger dynamics in dendritic spines,” Scientific reports, vol. 9, no. 1, pp. 1–18, 2019.

[117] L. A. Colgan and R. Yasuda, “Plasticity of dendritic spines: subcompartmentalization of signaling,” Annual review of physiology, vol. 76, pp. 365–385, 2014.

[118] R. Yasuda, “Biophysics of biochemical signaling in dendritic spines: implications in synaptic plasticity,” Biophysical journal, vol. 113, no. 10, pp. 2152–2159, 2017.

[119] E. Evans, H. Bowman, A. Leung, D. Needham, and D. Tirrell, “Biomembrane templates for nanoscale conduits and networks,” Science, vol. 273, no. 5277, pp. 933–935, 1996.

[120] D. Raucher and M. P. Sheetz, “Characteristics of a membrane reservoir buffering membrane tension,” Biophysical journal, vol. 77, no. 4, pp. 1992–2002, 1999.

[121] A. Roux, G. Cappello, J. Cartaud, J. Prost, B. Goud, and P. Bassereau, “A minimal system allowing tubulation with molecular motors pulling on giant liposomes,” Proceedings of the National Academy of Sciences, vol. 99, no. 8, pp. 5394–5399, 2002.

[122] T. A. Blanpied, D. B. Scott, and M. D. Ehlers, “Dynamics and regulation of clathrin coats at specialized endocytic zones of dendrites and spines,” Neuron, vol. 36, no. 3, pp. 435–449, 2002.

[123] G. L. Collingridge, J. T. Isaac, and Y. T. Wang, “Receptor trafficking and synaptic plasticity,” Nature Reviews Neuroscience, vol. 5, no. 12, pp. 952–962, 2004.

[124] C. T. Lee, J. G. Laughlin, N. A. de La Beaumelle, R. E. Amaro, J. A. McCammon, R. Ramamoorthi, M. Holst, and P. Rangamani, “3d mesh processing using gamer 2 to enable reaction-diffusion simulations in realistic cellular geometries,” PLoS computational biology, vol. 16, no. 4, p. e1007756, 2020.

[125] M. Akamatsu, R. Vasan, D. Serwas, M. A. Ferrin, P. Rangamani, and D. G. Drubin, “Principles of self-organization and load adaptation by the actin cytoskeleton during clathrin-mediated endocytosis,” Elife, vol. 9, p. e49840, 2020.

