## Supplementary material for "Mechanical principles governing dendritic spine shapes" for "Mechanical principles governing the shapes of dendritic spines"

**2 Supplementary figures** **11**

*

Table S1: Notations used in the model

| Notation | Description | Units |
|---|---|---|
| $E$ | Strain energy | $\mathrm{pN} \cdot \mathrm{nm}$ |
| $p$ | Pressure difference across the membrane | $\mathrm{pN{\cdot}nm^{-2}}$ |
| $\theta^\alpha$ | Surface parametrization | |
| $\sigma$ | Local energy per unit area | $\mathrm{pN \cdot nm^{-1}}$ |
| $\mathbf{r}$ | Position vector | |
| $\mathbf{n}$ | Normal vector to the membrane surface | unit vector |
| $\mathbf{a}_\alpha$ | Basis vector describing the tangent plane | |
| $\mathbf{a}^\alpha$ | Contravariant basis vector | |
| $\lambda$ | Tension, $-(W+\gamma)$ | $\mathrm{pN{\cdot}nm^{-1}}$ |
| $H$ | Mean curvature | $\mathrm{nm^{-1}}$ |
| $K$ | Gaussian curvature | $\mathrm{nm^{-2}}$ |
| $D$ | Curvature deviator | $\mathrm{nm^{-1}}$ |
| $D_m$ | Spontaneous deviatoric curvature | $\mathrm{nm^{-1}}$ |
| $\kappa$ | Bending modulus | $\mathrm{pN} \cdot \mathrm{nm}$ |
| $s$ | Arclength | nm |
| $\psi$ | Angle between $\mathbf{e}_r$ and $\mathbf{a}_s$ | |
| $r$ | Radial distance | nm |
| $z$ | Elevation from base plane | nm |
| $\mathbf{e}_r(\theta)$ | Radial basis vector | unit vector |
| $\mathbf{k}$ | Altitudinal basis vector | unit vector |
| $\mathbf{f}$ | Force density | $\mathrm{pN{\cdot}nm^{-2}}$ |
| $\mathrm{f_z}$ | Axial force density | $\mathrm{pN{\cdot}nm^{-2}}$ |
| $\mathrm{f_n}$ | Normal force density | $\mathrm{pN{\cdot}nm^{-2}}$ |
| $\mathrm{F_z}$ | Axial force | pN |
| $\kappa_\tau$ | Tangential curvature | $\mathrm{nm^{-1}}$ |
| $\kappa_\nu$ | Transverse curvature | $\mathrm{nm^{-1}}$ |
| $A$ | Total area of membrane | $\mathrm{nm^2}$ |
| $A_{\mathrm{force}}$ | Area of applied force | $\mathrm{nm^2}$ |
| $\boldsymbol{\gamma}$ | unit vector representing the one-dimensional orientation of a protein coat | |
| $\boldsymbol{\mu}$ | unit vector normal to $\boldsymbol{\gamma}$ and $\mathbf{n}$ | |
| $V$ | Confined volume by membrane area | $\mathrm{nm^3}$ |
| $A_{\mathrm{max}}$ | Maximum area of membrane | $\mathrm{nm^2}$ |
| $\lambda_0$ | Surface tension at boundary | $\mathrm{pN{\cdot}nm^{-1}}$ |
| $L$ | Membrane height | nm |
| $M$ | Shape equation variable | $\mathrm{nm^{-1}}$ |
| $x$ | Dimensionless radial distance | |
| $y$ | Dimensionless height | |
| $h$ | Dimensionless mean curvature | |
| $d$ | Dimensionless spontaneous deviatoric curvature | |
| $m$ | Dimensionless shape variable M | |
| $\lambda^*$ | Dimensionless tension | |
| $p^*$ | Dimensionless pressure | |
| $f^*$ | Dimensionless force density | |
| $\kappa^*$ | Dimensionless bending rigidity | |
| $\alpha^*$ | Dimensionless area | |

### 1 Model development

#### 1.1 Assumptions

- We treat the lipid bilayer as a continuous thin elastic shell assuming that the membrane thickness is negligible compared to the radii of membrane curvature [1]. This allows us to model the bending energy of the membrane using the modified version of the Helfrich–Canham energy including the effect of spatially varying deviatoric curvature to represent the induced anisotropic curvatures by periodic F-actin rings and other structures [2–7].
- We assume that the membrane is locally inextensible since the stretching modulus of the lipid bilayer is an order of magnitude larger than the membrane bending modulus [8]. We implemented this constraint using a Lagrange multiplier which can be interpreted as the tension [9–11]. We note that this membrane tension, in this study, is better interpreted as the effective contribution of the membrane in-plane stresses and membrane-cortex interactions [12].
- We assume that the time scales of mechanical forces are much faster than other events in dendritic spines allowing us to assume mechanical equilibrium and neglect inertia [9, 13]. This assumption is reasonable because the time scale of the equilibration of the mechanical forces is much smaller than the time scale of actin polymerization in dendritic spines [14].
- We assume that the force exerted by the actin cytoskeleton can be represented as work done on the membrane and do not include the molecular details of the actin network [13, 15–18]. Additionally, we assume that the periodic ring shaped structures of actin and related proteins such as $\beta$II spectrins, septics, and BAR-domain proteins can be represented using an anisotropic spontaneous curvature [4, 6, 7, 19].
- For ease of computation, we assume that the geometry of a dendritic spine is rotationally symmetric (see Fig. 1B) [13]. This assumption allows us to parametrize the whole surface by a single parameter which is the arclength.

#### 1.2 Membrane mechanics

At equilibrium, the total energy of the system ($E$) including the elastic storage energy of the membrane ($E_{\text{elastic}}$) and the work done by the applied forces by the actin filament ($W_{\text{force}}$) is given by [5, 15, 20, 21]

$$E = E_{\text{elastic}} - W_{\text{force}}, \tag{S1}$$

where

$$E_{\text{elastic}} = \int_{\omega} (\sigma(H, K, D; \theta^{\alpha}) + \lambda(\theta^{\alpha}))da - pV \quad \text{and} \tag{S2a}$$

$$W_{\text{force}} = \int_{\omega} \mathbf{f}(\theta^{\alpha}) \cdot (\mathbf{r} - \mathbf{r}_0)da. \tag{S2b}$$

Here $\omega$ is the total membrane surface area, $\sigma$ is the energy density, $\theta^{\alpha}$ denotes the surface coordinate where $\alpha \in \{1, 2\}$, $H$ is the mean curvature of the surface, $K$ is the Gaussian curvature, $D$ is the curvature deviator, $\lambda$ is the tension field which is the Lagrange multiplier associated with the local area constraint, $p$ is the transmembrane pressure that is the Lagrange multiplier associated with the volume constraint, $V$ is the enclosed volume, $\mathbf{f}$ is the applied force per unit area, $\mathbf{r}$ is the position vector in the current configuration, and $\mathbf{r}_0$ is the position vector in the reference frame. We can write the variation of the total free energy of the system as [15, 20]

$$\dot{E} = \dot{E}_{\text{elastic}} - \dot{W}_{\text{force}}, \tag{S3}$$

where

$$\dot{E}_{\text{elastic}} = \int_{\omega} \dot{\sigma}\, da + \int_{\omega} (\sigma + \lambda)(\dot{J}/J)\, da - p\dot{V} \quad \text{and} \tag{S4a}$$

$$\dot{W}_{\text{force}} = \int_{\omega} \mathbf{f}(\theta^{\alpha}) \cdot \mathbf{u}\, da, \tag{S4b}$$

where $J = \sqrt{a/A}$ is the local areal stretch due to mapping from a reference frame ($A$) to the actual surface ($a$), and $\mathbf{u}$ is the virtual displacement of the surface given by

$$\mathbf{u}(\theta^{\alpha}) = \frac{\partial}{\partial \epsilon} \mathbf{r}(\theta^{\alpha}, \epsilon)|_{\epsilon=0} = \dot{\mathbf{r}}. \tag{S5}$$

Minimization of the energy in Eq. S3 by usage of the variational approach gives the governing shape equation and the incompressibility condition in a heterogeneous membrane as

$$\begin{aligned} p + \mathbf{f} \cdot \mathbf{n} = {} & \Delta(\frac{1}{2}\sigma_H) + (\sigma_K)_{;\alpha\beta}\tilde{b}^{\alpha\beta} + \sigma_H(2H^2 - K) + 2H(K\sigma_K - \sigma) - 2\lambda H \\ & + \frac{1}{2}[\sigma_D(\gamma^{\alpha}\gamma^{\beta} - \mu^{\alpha}\mu^{\beta})]_{;\beta\alpha} + \frac{1}{2}\sigma_D(\gamma^{\alpha}\gamma^{\beta} - \mu^{\alpha}\mu^{\beta}) b_{\alpha\eta} b^{\eta}_{\beta}, \end{aligned} \tag{S6}$$

and

$$(\frac{\partial \sigma}{\partial \theta^{\alpha}_{|exp}} + \lambda_{,\alpha} + \sigma_D[b_{\alpha\beta}(\gamma^{\alpha}\gamma^{\beta})_{;\eta}]) a^{\beta\alpha} = \mathbf{f} \cdot \mathbf{a}_s, \tag{S7}$$

where $\Delta(\cdot)$ is the surface Laplacian, $\mathbf{n}$ is the normal vector to the surface, $\mathbf{a}_s$ is a tangent vector on the surface (we will define it in the next section for axisymmetric coordinates), $a^{\alpha\beta}$ is the dual metric, $b_{\alpha\beta}$ are the coefficients of the second fundamental form, $b^{\alpha}_{\beta}$ are the mixed components of the curvature, $\tilde{b}^{\alpha\beta}$ is the co-factor of the curvature tensor, $(.)_{;\alpha}$ is the covariant derivative, $(.)_{,\alpha}$ is the partial derivative, and $()_{|exp}$ denotes the explicit derivative with respect to coordinate $\theta^{\alpha}$. Also, $\gamma^{\alpha}$ and $\mu^{\alpha}$ are the projections of $\boldsymbol{\gamma}$ and $\boldsymbol{\mu}$ along the tangential vectors given by [20]

$$\begin{aligned} \gamma^{\alpha} &= \boldsymbol{\gamma} \cdot \mathbf{a}^{\alpha} \\ \mu^{\alpha} &= \boldsymbol{\mu} \cdot \mathbf{a}^{\alpha}, \end{aligned} \tag{S8}$$

where $\mathbf{a}^{\alpha}$ is the contravariant basis vectors, $\boldsymbol{\gamma}$ is a unit vector representing the orientation of a one-dimensional curve on the surface which is tangential to the protein coat, and $\boldsymbol{\mu}$ is a unit vector defined as

$$\boldsymbol{\mu} = \mathbf{n} \times \boldsymbol{\gamma}. \tag{S9}$$

##### 1.3 Helfrich energy including deviatoric curvature

We modeled the combined effects of BAR domain proteins and periodic F-actin by deviatoric curvature using the modified version of Helfrich energy that includes deviatoric curvature $D(\theta^{\alpha})$ [2–7, 22] given as

$$\sigma(H,K,D;\theta^\alpha) = \kappa H^2 + \kappa(D - D_m(\theta^\alpha))^2, \tag{S10}$$

where $\kappa$ is the bending modulus and $D_m$ is the spontaneous (intrinsic) deviatoric curvature [20, 23]. Substituting this form of energy density (Eq. S10) in Eqs. S6 and S7 gives

$$\begin{aligned}\underbrace{-\kappa\Big[2H(D-D_m)^2 - \Big((D-D_m)(\gamma^\alpha\gamma^\beta - \mu^\alpha\mu^\beta)\Big)_{;\beta\alpha} - (D-D_m)(\gamma^\alpha\gamma^\beta - \mu^\alpha\mu^\beta)b_{\alpha\eta}b^\eta_\beta\Big]}_{\text{Induced anistropic curvature effects}} \\ + \underbrace{\kappa\Delta H + 2\kappa H(H^2-K)}_{\text{Elastic effects}} = \underbrace{p + 2\lambda H}_{\text{Capillary effects}} + \underbrace{\mathbf{f}\cdot\mathbf{n}}_{\text{Force due to actin}},\end{aligned} \tag{S11}$$

and

$$\underbrace{\lambda_{,\alpha}}_{\substack{\text{Tension}\\ \text{variation}}} = \underbrace{2\kappa(D-D_m)\frac{\partial D_m}{\partial\theta^\alpha} + 2\kappa(D-D_m)b_{\alpha\beta}(\gamma^\alpha\gamma^\beta)_{;\eta}}_{\text{Anisotropic curvature induced variation}} - \underbrace{\mathbf{f}\cdot\mathbf{a_s}}_{\substack{\text{Force-induced}\\ \text{variation}}}. \tag{S12}$$

It should be mentioned that in the modified version of Helfrich energy ( Eq. S10), we assumed that the induced isotropic spontaneous curvature ($C$) by BAR domain proteins or periodic F-actin is negligible and we ignored the effect of the spontaneous curvature ($C = 0$).

#### 1.4 Governing equations in axisymmetric coordinates

##### 1.4.1 Axisymmetric coordinates

We parameterize a surface of revolution with respect to the z axis (Fig. 1B) in the coordinate basis $(\mathbf{e}_r, \mathbf{e}_\theta, \mathbf{k})$ by

$$\mathbf{r}(s,\theta) = r(s)\mathbf{e}_r(\theta) + z(s)\mathbf{k}, \tag{S13}$$

where $s$ is the arclength along the curve, $r(s)$ is the radial distance from the axis of rotation, and $z(s)$ is the elevation from the reference plane. Since $(dr/ds)^2 + (dz/ds)^2 = 1$, we can define $\psi$ (the angle made by the tangent with respect to the horizontal) such that the normal and tangent vectors are given by

$$\mathbf{n} = -\sin\psi\mathbf{e}_r(\theta) + \cos\psi\mathbf{k} \quad \text{and} \quad \mathbf{a}_s = \cos\psi\mathbf{e}_r(\theta) + \sin\psi\mathbf{k}. \tag{S14}$$

Following this we have

$$r'(s) = \cos(\psi), \tag{S15a}$$
$$z'(s) = \sin(\psi), \tag{S15b}$$

where $(.)' = \frac{d(.)}{ds}$. We can now write the tangential ($\kappa_\nu$) and transverse ($\kappa_\tau$) principal curvatures as

$$\kappa_\nu = \psi', \quad \kappa_\tau = r^{-1}\sin\psi, \tag{S16}$$

and the mean curvature ($H$), Gaussian curvature ($K$), and the curvature deviator ($D$) as

$$\begin{aligned} H &= \frac{1}{2}(\kappa_\nu + \kappa_\tau) = \frac{1}{2}(\psi' + r^{-1}\sin\psi), \\ K &= \kappa_\tau\kappa_\nu = \frac{\psi'\sin\psi}{r}, \\ D &= \frac{1}{2}(\kappa_\tau - \kappa_\nu) = \frac{1}{2}(r^{-1}\sin\psi - \psi') = r^{-1}\sin\psi - H. \end{aligned} \tag{S17}$$

##### 1.4.2 Equilibrium equations

In order to simplify the governing shape and incompressibility equations, we define $M$ as

$$M = \frac{1}{2\kappa} r[(\sigma_H)' - (\sigma_D)'], \tag{S18}$$

which allows us to simplify the shape equation (Eq. S11) and the inextensibility condition (Eq. S12) as a system of first order differential equations given by

$$\begin{aligned}
r' = \cos\psi, \quad z' = \sin\psi, \quad r\psi' = 2rH - \sin\psi, \\
2rH' = M - D_m' + 2H\cos(\psi) - \frac{2\cos(\psi)\sin(\psi)}{r}, \\
\frac{M'}{r} = \frac{p}{\kappa} + \frac{\mathbf{f}\cdot\mathbf{n}}{\kappa} + 2H\Big[H^2 + \frac{\lambda}{\kappa} + (\frac{\sin(\psi)}{r} - H - D_m)^2 - 2(\frac{\sin(\psi)}{r} - H - D_m)(\frac{\sin(\psi)}{r} - H)\Big] \\
-2H\left[H^2 + \left(H - r^{-1}\sin\psi\right)^2\right] - 2\frac{\cos(\psi)}{r}\Big[\frac{H\cos(\psi)}{r} - \frac{\sin(\psi)\cos(\psi)}{r^2} - \frac{D_m'}{2} - \frac{M}{2r}\Big], \\
\lambda' = -2\kappa(\frac{\sin(\psi)}{r} - H - D_m)D_m' - \mathbf{f}\cdot\mathbf{a_s}.
\end{aligned} \tag{S19}$$

In axisymmetric coordinates, the total area of the manifold (A) can be expressed in term of arclength as

$$A(s) = 2\pi\int_0^s r(t)dt \quad \rightarrow \quad \frac{dA}{ds} = 2\pi r. \tag{S20}$$

This allows us to write the governing differential equations (Eq. S19) in terms of the derivative of area given as

$$\begin{aligned}
2\pi r\dot{r} = \cos\psi, \quad 2\pi r\dot{z} = \sin\psi, \quad r^2\dot{\psi} = 2rH - \sin\psi, \\
2\pi r^2\dot{H} = M - 2\pi r\dot{D}_m + 2H\cos(\psi) - \frac{2\cos(\psi)\sin(\psi)}{r}, \\
2\pi\dot{M} = \frac{p}{\kappa} + \frac{\mathbf{f}\cdot\mathbf{n}}{\kappa} + 2H\Big[H^2 + \frac{\lambda}{\kappa} + (\frac{\sin(\psi)}{r} - H - D_m)^2 - 2(\frac{\sin(\psi)}{r} - H - D_m)(\frac{\sin(\psi)}{r} - H)\Big] \\
-2H\left[H^2 + \left(H - r^{-1}\sin\psi\right)^2\right] - 2\frac{\cos(\psi)}{r}\Big[\frac{H\cos(\psi)}{r} - \frac{\sin(\psi)\cos(\psi)}{r^2} - \frac{\dot{D}_m}{2} - \frac{M}{2r}\Big] \\
2\pi r\dot{\lambda} = -4\pi r\kappa(\frac{\sin(\psi)}{r} - H - D_m)\dot{D}_m - \mathbf{f}\cdot\mathbf{a_s},
\end{aligned} \tag{S21}$$

where $\dot{(.)} = \frac{d(.)}{dA}$. In order to solve the coupled partial differential equations in Eq. S21, we impose six boundary conditions as follow:

$$\begin{aligned}
r(0^+) = 0, \quad M(0^+) = 0, \quad \psi(0^+) = 0, \\
z(A_{\text{max}}) = 0, \quad \psi(A_{\text{max}}) = 0, \quad \lambda(A_{\text{max}}) = \lambda_0,
\end{aligned} \tag{S22}$$

where $\lambda_0$ is the tension at the far field boundary. We can non-dimensionalize the system of equations in Eq. S21, using two constants $R_0$ and $\kappa_0$, defining the non-dimensional parameters as

$$\begin{aligned}
\alpha = \frac{A}{2\pi R_0^2}, \quad x = \frac{r}{R_0}, \quad y = \frac{z}{R_0}, \quad h = HR_0, \quad d = D_m R_0, \quad m = MR_0, \\
\lambda^* = \frac{\lambda R_0^2}{\kappa_0}, \quad p^* = \frac{pR_0^3}{\kappa_0}, \quad f^* = \frac{fR_0^3}{\kappa_0}, \quad \kappa^* = \frac{\kappa}{\kappa_0}.
\end{aligned} \tag{S23}$$

Using these non-dimensional parameters and substituting into the system of partial differential equations (Eq. S21), we get

$$\begin{aligned} x\dot{x} &= \cos\psi, \quad x\dot{x} = \sin\psi, \quad x^2\dot{\psi} = 2xh - \sin\psi, \\ x^2\dot{h} &= m - x\dot{d} + 2h\cos(\psi) - \frac{2\cos(\psi)\sin(\psi)}{x}, \\ \dot{m} = \frac{p^*}{\kappa^*} + \frac{\mathbf{f}^*\cdot\mathbf{n}}{\kappa^*} &+ 2h\Big[h^2 + \frac{\lambda^*}{\kappa^*} + (\frac{\sin(\psi)}{x} - h - d)^2 - 2(\frac{\sin(\psi)}{x} - h - d)(\frac{\sin(\psi)}{x} - h)\Big] \\ -2h\left[h^2 + \left(h - x^{-1}\sin\psi\right)^2\right] &- 2\frac{\cos(\psi)}{x}\Big[\frac{h\cos(\psi)}{x} - \frac{\sin(\psi)\cos(\psi)}{x^2} - \frac{\dot{d}}{2} - \frac{m}{2x}\Big], \\ \dot{\lambda} &= -2\kappa^*(\frac{\sin(\psi)}{x} - h - d)\dot{d} - \mathbf{f}^*\cdot\mathbf{a_s}, \end{aligned} \tag{S24}$$

and the boundary conditions, (Eq. S22), become

$$\begin{aligned} x(0^+) = 0, \quad m(0^+) = 0, \quad \psi(0^+) = 0, \\ y(\alpha_{\text{max}}) = 0, \quad \psi(\alpha_{\text{max}}) = 0, \quad \lambda^*(\alpha_{\text{max}}) = \lambda_0^*. \end{aligned} \tag{S25}$$

##### 1.4.3 Numerical implementation

In order to solve the system of differential equations (Eq. S24) along with the boundary conditions (Eq. S25), we used 'bvp4c,' a boundary value problem solver in MATLAB. In all simulations, we fixed the total area of the membrane as $A = 8\pi\mu\text{m}^2$ and set the transmembrane pressure to be zero ($p = 0$) to focus mainly on the mechanism of membrane-actin interactions in dendritic spine formation. The mesh points on the domain were chosen such that starting from $A = 0^+$, the mesh size is very small and then increases moving toward the far field boundary $A = A_{\text{max}}$. To have a sharp but smooth transition at the boundaries of the applied forces, we prescribed the forces using a hyperbolic tangent function given as

$$f = \frac{f_0}{2}[\tanh(g(A - A_{\text{force}}))], \tag{S26}$$

where $g$ is a constant and $A_{\text{force}}$ represents the area of the applied forces by actin filaments. Additionally, to get the tubular protrusions from a flat membrane, we prescribed the height of the protrusion as an extra boundary condition ($z(0^+) = z_p$) and calculated the magnitude of the applied force as an unknown parameter.

#### 1.5 Analytical solutions

##### 1.5.1 Analytical estimation for filopodia-shaped spines with an axial force

Let us consider a long tubular filopodium with radius r and height L has been pulled from a flat membrane with a axial force $\text{F}_\text{z}$ (Fig. S1A). Assuming that $\text{L} \gg \text{r}$, in the absence of spontaneous deviatoric curvature ($D_m = 0$) and pressure ($p = 0$), the total free energy of the system (Eq. S1) can be written as

$$E_{\text{filopodium}} = \int_\omega (\kappa H^2 + \kappa D^2 + \lambda)da - \text{F}_\text{z}\text{L}. \tag{S27}$$

For a tubular membrane (ignoring the spherical cap), $H = D = \frac{1}{2\text{r}}$, $\int da = 2\pi rL$, and thus the total energy can be simplified as

$$E_{\text{filopodium}} = (\frac{\kappa}{2\text{r}^2} + \lambda)2\pi\text{rL} - \text{F}_\text{z}\text{L}. \tag{S28}$$

Now, we can find the equilibrium radius of the tube ($r$) and the corresponding force ($F_z$) by taking $\partial E_{\text{filopodium}}/\partial \mathrm{r} = 0$ and $\partial E_{\text{filopodium}}/\partial \mathrm{L} = 0$ and solving for the radius and force as [24–26]

$$\mathrm{r} = \sqrt{\frac{\kappa}{2\lambda}} \quad \text{and} \quad \mathrm{F_z} = 2\pi\sqrt{2\lambda\kappa}. \tag{S29}$$

Based on the Eq. S29, the diameter of the filopodium and the magnitude of the applied forces depend on the tension and the bending rigidity. For example, for a fixed bending rigidity, a large force is required to bend a stiff membrane (large tension) and form a narrow filopodium. Interestingly, in contrast to the stubby spine (Fig.2C), the magnitude of force to form a tubular filopodium is independent of the length of protrusion. In Eq. S29, we can also find the tension based on the radius of the tubule and rewrite the axial force as

$$\mathrm{F_z} = \frac{2\pi\kappa}{\mathrm{r}}, \tag{S30}$$

which indicates that to form a narrower filopodium, a larger axial force is required.

##### 1.5.2 Analytical estimation for thin-shaped spines with a uniform normal force density

Let us consider an idealized geometry of a thin-shaped spine as a sphere with radius R that is connected to a cylinder with radius $r$ and height $l$ (Fig. S1B). Considering the case that a uniform normal force density, $\mathrm{f_n}$, is applied all along the sphere and ignoring the interface between the sphere and the cylinder, we can write the total energy of the system as $E_{\text{thin}} = E_{\text{sphere}} + E_{\text{cylinder}}$. For the sphere, we know that $H = 1/\mathrm{R}$, $D = 0$, and the total surface area is $A_{\text{sphere}} = 4\pi\mathrm{R}^2$. Considering the axial displacement from a flat membrane, we can write the free energy of the spherical part of the thin-shaped spine as (assuming $p = 0$ and $D_m = 0$)

$$E_{\text{sphere}} = (\frac{\kappa}{\mathrm{R}^2} + \lambda)4\pi\mathrm{R}^2 - \int -\mathrm{f_n}\cos(\phi)\big(\mathrm{L} + \mathrm{R} - \mathrm{R}\cos(\phi)\big)da, \tag{S31}$$

where $da$ is the area element which can be written as $\int da = \int_0^\pi 2\pi\mathrm{R}^2\sin(\phi)d\phi$. Eq. S31 gives

$$E_{\text{sphere}} = (\frac{\kappa}{\mathrm{R}^2} + \lambda)4\pi R^2 - \frac{4\pi}{3}\mathrm{R}^3\mathrm{f_n}. \tag{S32}$$

For the cylindrical part of the thin-shaped spine, based on the previous section for the filopodia, we know that $H = D = 1/2\mathrm{r}$ and $\mathrm{r} = \sqrt{\kappa/2\lambda}$. Substituting these terms in the energy, we get

$$E_{\text{cylinder}} = 2\pi\sqrt{2\lambda\kappa}l. \tag{S33}$$

Using Eqs. S32 and S33, we can write the total energy of the system as

$$E_{\text{thin}} = (\frac{\kappa}{\mathrm{R}^2} + \lambda)4\pi\mathrm{R}^2 - \frac{4\pi}{3}\mathrm{R}^3\mathrm{f_n} + 2\pi\sqrt{2\lambda\kappa}l. \tag{S34}$$

Taking $\partial E_{\text{thin}}/\partial\mathrm{R} = 0$, we obtain

$$\mathrm{f_n} = \frac{2\lambda}{\mathrm{R}}. \tag{S35}$$

Eq. S35 resembles the Young-Laplace equation where, instead of pressure (normal force density) as a global parameter, $f_n$ is a local normal force density. To find the total normal force ($F_n$), we need to multiply the force density in Eq. S35 by the total area of the applied force which is the area of sphere ($A_{force} = A_{sphere}$). This gives us

$$F_n = f_n \times \mathit{A}_{force} = f_n \times 4\pi R^2 = 8\pi R\lambda. \tag{S36}$$

In our simulation, we are prescribing the area of the applied normal force ($A_{force}$) based on the area of the spine head ($A_{spine\text{-}head}$). Assuming that the spine head has a spherical shape in the area of the applied force, we can find the radius of the sphere based on the area of spine head as

$$A_{force} = A_{spine\text{-}head} = A_{sphere} = 4\pi R^2 \rightarrow R = \sqrt{\frac{A_{spine\text{-}head}}{4\pi}}. \tag{S37}$$

Substituting Eq. S37 into Eq. S36, we have

$$F_n = 2\lambda\sqrt{4\pi A_{spine\text{-}head}}. \tag{S38}$$

Based on Eq. S38, if we know the area of the spine head and the tension, we can estimate the required normal force to form a thin-shaped spine. For example, according to Eq. S38, a larger normal force is required to form a thin-shaped spine with a larger head. Similar to the formation of a filopodium (Eq. S29), the magnitude of the normal force in Eq. S38 is also independent of the length of the spine.

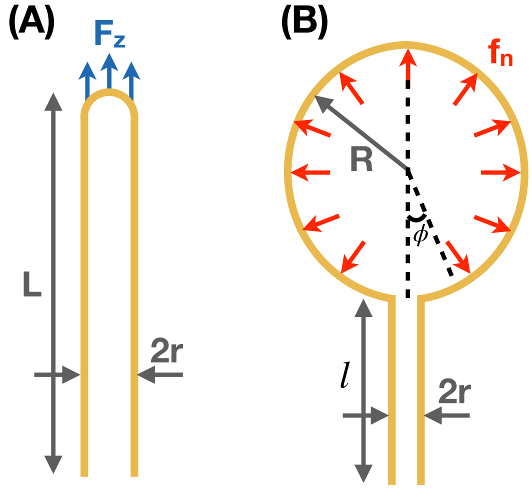

Figure S1: (A) Schematic of a long filopodium with radius r and height L formed with an axial force $F_z$ applied along the prescribed area shown in blue. (B) An idealized geometry of a thin shaped spine. A sphere with radius R is connected to a cylinder with radius r and height $l$. A uniform normal force density $f_n$ is applied all along the sphere ($A_{force} = A_{sphere} = A_{spine\text{-}head}$).

Additionally, since the spine neck has a tubular shape, we can relate the tension to the radius of spine neck (Eq. S29) and rewrite the normal force as

$$F_n = \frac{\kappa}{r^2}\sqrt{4\pi A_{spine\text{-}head}}, \tag{S39}$$

where r is the radius of the spine neck (Fig. S1B). Comparison of Eq. S30 and Eq. S39 shows that the required axial force to form a tubular filopodium is proportional to $1/r$ while the normal force to form a thin-shaped spine is proportional to $1/r^2$. This suggests that decreasing the neck diameter of a thin-shaped spines is harder and needs a larger magnitude of a force compared to the filopodial-shaped spines.

##### 1.5.3 Spontaneous deviatoric curvature and the radius of the spines neck

Let us consider a tubular membrane with radius r and height L that has been pulled from a flat membrane with an axial force $\mathrm{F_z}$ and a spontaneous deviatoric curvature $D_m$ along the neck region (Fig. S2A). Ignoring the spherical cap, the total free energy of the system (Eq. S1) can be written as

$$E_{\text{tube}} = \Big(\frac{\kappa}{4\mathrm{r}^2} + \lambda + \kappa(\frac{1}{2\mathrm{r}} - D_m)^2\Big)2\pi\mathrm{rL} - \mathrm{F_z}l. \tag{S40}$$

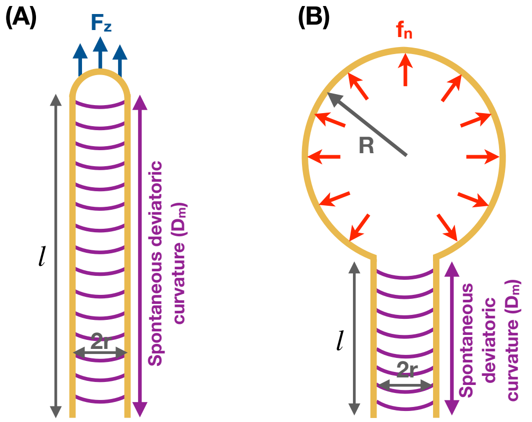

Figure S2: Schematic of (A) a tubular membrane with an axial force ($\mathrm{F_z}$) and spontaneous deviatoric curvature ($D_m$) along the neck with length $l$ and (B) an idealized geometry of a thin-shaped spine with normal force density ($\mathrm{f_n}$) along the head and spontaneous deviatoric curvature ($D_m$) along the neck.

To find the equilibrium radius of the tube ($r$) and the corresponding force ($\mathrm{F_z}$), we take $\partial E_{\text{tube}}/\partial \mathrm{r} = 0$ and $\partial E_{\text{tube}}/\partial l = 0$ and solve for the radius and force as

$$\mathrm{r} = \sqrt{\frac{\kappa}{2(\lambda + \kappa D_m^2)}} \quad \text{and} \quad \mathrm{F_z} = 2\pi\Big(\sqrt{2\kappa(\lambda + \kappa D_m^2)} - \kappa D_m\Big), \tag{S41}$$

which reduces to Eq. S29 for zero spontaneous deviatoric curvature ($D_m = 0$). Based on Eq. S41, we can see that the radius of a tubule decreases with increasing magnitude of spontaneous deviatoric curvature (Fig. 5A). This is consistent with the previous study by Walani et al. where they showed that a tubular membrane gets narrower with increasing strength of spontaneous deviatoric curvature [20]. In Fig. 5C, we plotted the magnitude of the axial force as a function of tension and spontaneous deviatoric curvature. As can be seen, axial force has a local minimum shown by the red dashed line. By taking $\partial F_z/\partial D = 0$, we can find the relationship between the tension and spontaneous deviatoric curvature along the red line (Fig. 5C) is

$$\lambda = \kappa D_m^2, \tag{S42}$$

and the magnitude of local minimum force is given by

$$\mathrm{F_{z,min}} = 2\pi\kappa D_m, \tag{S43}$$

which linearly depends on $D_m$. In addition, we can use the radius-tension relationship in Eq. S41 to revise the normal force equation for the formation of thin-shaped spines (Eq. S39) to include the effect of spontaneous deviatoric curvature and get

$$F_n = 2\kappa(\frac{1}{2r^2} - D_m^2)\sqrt{4\pi A_{\text{spine-head}}}, \quad (S44)$$

which indicates that by adding the effect of spontaneous deviatoric curvature, the magnitude of normal force needed to stabilize a thin-shaped spine decreases. Theoretically, if $D_m = \frac{1}{\sqrt{2}r_{\text{neck}}}$, a thin-shaped spine can have a stable geometry without any actin-mediated forces.

#### 2 Supplementary figures

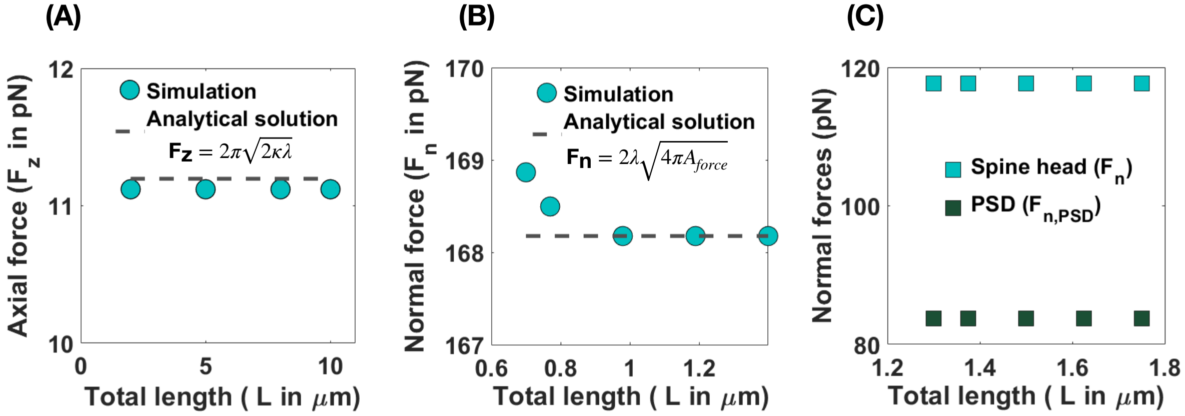

Figure S3: The magnitude of axial and normal forces that are required to form (A) filopodia, (B) thin, and (C) mushroom-shaped spines are independent of the length of the spines.

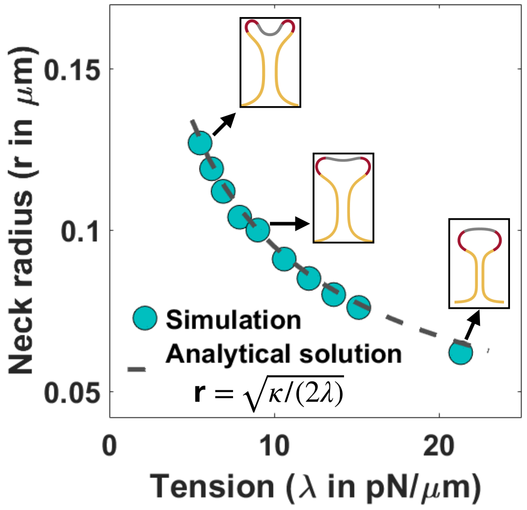

Figure S4: (A) Neck radius of a mushroom-shaped spine as a function of tension for area of PSD/area of head = 0.2 ($r = \sqrt{\kappa/(2\lambda)}$) [24]. Three different shapes of mushroom-shaped spines are shown for low, intermediate, and high tension. With increasing magnitude of tension, the mushroom-shaped spine flattens.

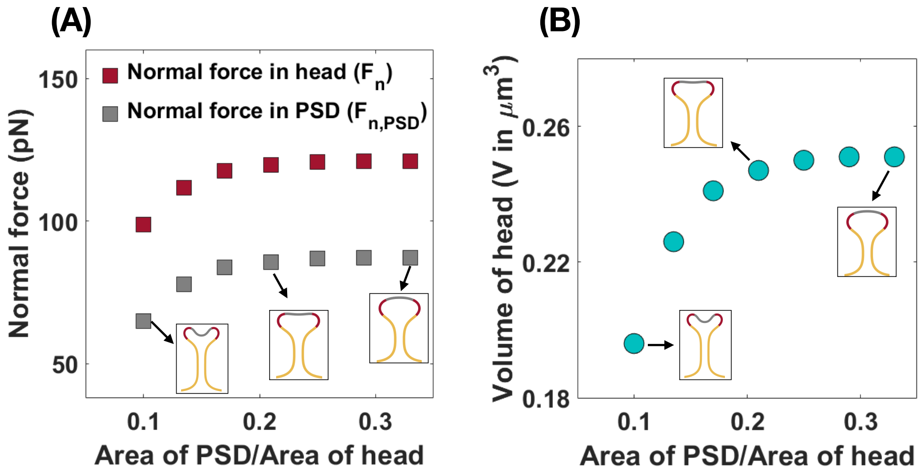

Figure S5: The area of PSD with respect to the area of the spine head characterizes the normal forces that are required to form a mushroom spine. (A) The magnitude of normal forces in the spine head (red squares) and in the PSD (gray squares) increases with increasing ratio of PSD area to head area. The mushroom-shaped spine flattens with increasing ratio of PSD area to head area. (B) A larger mushroom-shaped spine (larger head volume) forms with increasing ratio of PSD area to head area.

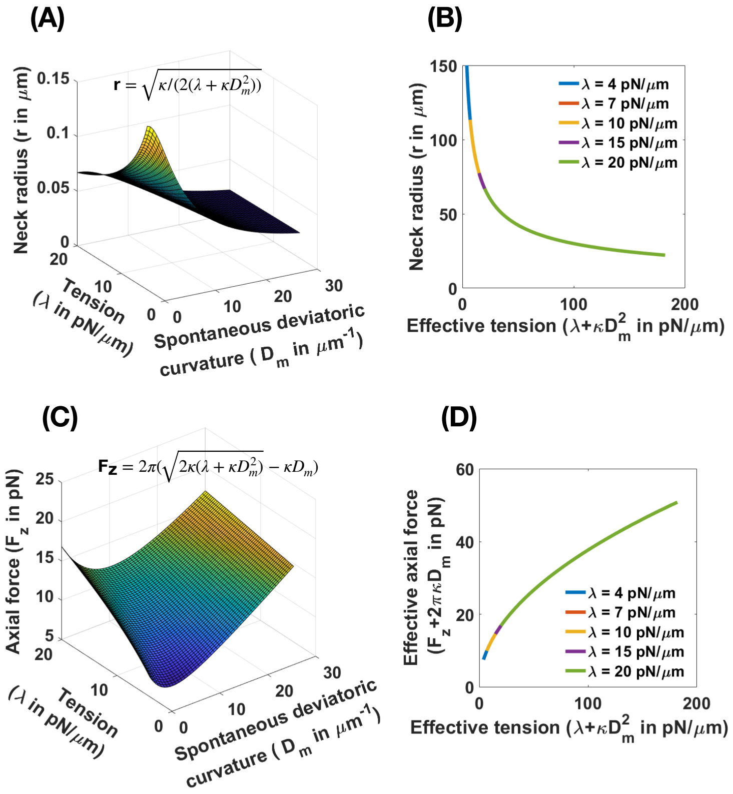

Figure S6: (A) Analytical solution for the neck radius of a tubular membrane as functions of spontaneous deviatoric curvature and tension ($\mathrm{r} = \sqrt{\kappa/(2(\lambda+\kappa \mathrm{D}_\mathrm{m}^2))}$, Eq. S41). (B) The neck radius in panel A collapses onto a single curve as a function of effective tension. (C) Analytical solution for the magnitude of an axial force needed to maintain a tubular protrusion as functions of spontaneous deviatoric curvature and tension ($\mathrm{F_z} = 2\pi(\sqrt{2\kappa(\lambda+\kappa \mathrm{D}_\mathrm{m}^2)} - \kappa \mathrm{D_m})$, Eq. S41). (D) The effective axial force in panel C collapses onto a single curve as a function of effective tension.

Table S2: Energy components and total energy for three different mechanisms of thin spine formation

| | Bending energy (pN.$\mu$m) $\int \kappa H^2 da$ | Bending energy due to deviatoric curvature (pN.$\mu$m) $\int \kappa (D - D_m)^2 da$ | Work done by force (pN.$\mu$m) $\int \mathbf{f} \cdot (\mathbf{r} - \mathbf{r}_0) da$ | Work done by tension (pN.$\mu$m) $\int \lambda da$ | Total energy (pN.$\mu$m) |
|---|---|---|---|---|---|
| Uniform normal force density along head | 6.2 | 6.2 | -27.64 | 904.8 | 899.5 |
| Uniform normal force density along head and spontaneous deviatoric curvature along neck | 5.8 | 1.75 | -16.35 | 254.34 | 245.5 |
| Uniform axial force density along head and spontaneous deviatoric curvature along neck | 5.28 | 1 | -9.1 | 247.1 | 244.3 |

Table S3: Energy components and total energy for three different mechanisms of mushroom spine formation

| | Bending energy (pN.$\mu$m) $\int \kappa H^2 da$ | Bending energy due to deviatoric curvature (pN.$\mu$m) $\int \kappa (D - D_m)^2 da$ | Work done by force (pN.$\mu$m) $\int \mathbf{f} \cdot (\mathbf{r} - \mathbf{r}_0) da$ | Work done by tension (pN.$\mu$m) $\int \lambda da$ | Total energy (pN.$\mu$m) |
|---|---|---|---|---|---|
| Non-uniform normal force density along head | 4.92 | 4.92 | -18.65 | 226.2 | 217.4 |
| Non-uniform normal force density along head and spontaneous deviatoric curvature along neck | 513 | 4.29 | -17.43 | 139.2 | 131 |

### References

1. H. Deuling and W. Helfrich, "Red blood cell shapes as explained on the basis of curvature elasticity," *Biophysical journal*, vol. 16, no. 8, pp. 861–868, 1976.

2. W. Helfrich, "Elastic properties of lipid bilayers: theory and possible experiments," *Zeitschrift fur Naturforschung C*, vol. 28, no. 11-12, pp. 693–703, 1973.

3. P. B. Canham, "The minimum energy of bending as a possible explanation of the biconcave shape of the human red blood cell," *Journal of theoretical biology*, vol. 26, no. 1, pp. 61–81, 1970.

4. A. Iglič, H. Hägerstrand, P. Veranič, A. Plemenitaš, and V. Kralj-Iglič, "Curvature-induced accumulation of anisotropic membrane components and raft formation in cylindrical membrane protrusions," *Journal of Theoretical Biology*, vol. 240, no. 3, pp. 368–373, 2006.

5. H. Alimohamadi and P. Rangamani, "Modeling membrane curvature generation due to membrane–protein interactions," *Biomolecules*, vol. 8, no. 4, p. 120, 2018.

6. V. Kralj-Iglič, V. Heinrich, S. Svetina, and B. Žekš, "Free energy of closed membrane with anisotropic inclusions," *The European Physical Journal B-Condensed Matter and Complex Systems*, vol. 10, no. 1, pp. 5–8, 1999.

7. V. Kralj-Iglič, S. Svetina, and B. Žekž, "Shapes of bilayer vesicles with membrane embedded molecules," *European biophysics journal*, vol. 24, no. 5, pp. 311–321, 1996.

8. W. Rawicz, K. Olbrich, T. McIntosh, D. Needham, and E. Evans, "Effect of chain length and unsaturation on elasticity of lipid bilayers," *Biophysical journal*, vol. 79, no. 1, pp. 328–339, 2000.

9. D. Steigmann, "Fluid films with curvature elasticity," *Archive for Rational Mechanics and Analysis*, vol. 150, no. 2, pp. 127–152, 1999.

10. P. Rangamani, A. Agrawal, K. K. Mandadapu, G. Oster, and D. J. Steigmann, "Interaction between surface shape and intra-surface viscous flow on lipid membranes," *Biomechanics and modeling in mechanobiology*, pp. 1–13, 2013.

11. H. Alimohamadi, B. Ovryn, and P. Rangamani, “Modeling membrane nanotube morphology: the role of heterogeneity in composition and material properties,” *Scientific reports*, vol. 10, no. 1, pp. 1–15, 2020.

12. A. Diz-Muñoz, D. A. Fletcher, and O. D. Weiner, “Use the force: membrane tension as an organizer of cell shape and motility,” *Trends in cell biology*, vol. 23, no. 2, pp. 47–53, 2013.

13. C. Miermans, R. Kusters, C. Hoogenraad, and C. Storm, “Biophysical model of the role of actin remodeling on dendritic spine morphology,” *PloS one*, vol. 12, no. 2, p. e0170113, 2017.

14. J. Weichsel and P. L. Geissler, “The more the tubular: Dynamic bundling of actin filaments for membrane tube formation,” *PLoS computational biology*, vol. 12, no. 7, p. e1004982, 2016.

15. N. Walani, J. Torres, and A. Agrawal, “Endocytic proteins drive vesicle growth via instability in high membrane tension environment,” *Proceedings of the National Academy of Sciences*, vol. 112, no. 12, pp. E1423–E1432, 2015.

16. E. Atilgan, D. Wirtz, and S. X. Sun, “Mechanics and dynamics of actin-driven thin membrane protrusions,” *Biophysical journal*, vol. 90, no. 1, pp. 65–76, 2006.

17. J. E. Hassinger, G. Oster, D. G. Drubin, and P. Rangamani, “Design principles for robust vesiculation in clathrin-mediated endocytosis,” *Proceedings of the National Academy of Sciences*, vol. 114, no. 7, pp. E1118–E1127, 2017.

18. H. Alimohamadi, A. S. Smith, R. B. Nowak, V. M. Fowler, and P. Rangamani, “Non-uniform distribution of myosin-mediated forces governs red blood cell membrane curvature through tension modulation,” *PLOS Computational Biology*, vol. 16, no. 5, p. e1007890, 2020.

19. A. Iglič, B. Babnik, K. Bohinc, M. Fošnarič, H. Hägerstrand, and V. Kralj-Iglič, “On the role of anisotropy of membrane constituents in formation of a membrane neck during budding of a multicomponent membrane,” *Journal of biomechanics*, vol. 40, no. 3, pp. 579–585, 2007.

20. N. Walani, J. Torres, and A. Agrawal, “Anisotropic spontaneous curvatures in lipid membranes,” *Physical Review E*, vol. 89, no. 6, p. 062715, 2014.

21. M. Lokar, D. Kabaso, N. Resnik, K. Sepčić, V. Kralj-Iglič, P. Veranič, R. Zorec, and A. Iglič, “The role of cholesterol-sphingomyelin membrane nanodomains in the stability of intercellular membrane nanotubes,” *International journal of nanomedicine*, vol. 7, p. 1891, 2012.

22. A. Iglič, B. Babnik, U. Gimsa, and V. Kralj-Iglič, “On the role of membrane anisotropy in the beading transition of undulated tubular membrane structures,” *Journal of Physics A: Mathematical and General*, vol. 38, no. 40, p. 8527, 2005.

23. H. Alimohamadi, R. Vasan, J. E. Hassinger, J. C. Stachowiak, and P. Rangamani, “The role of traction in membrane curvature generation,” *Molecular biology of the cell*, vol. 29, no. 16, pp. 2024–2035, 2018.

24. I. Derényi, F. Jülicher, and J. Prost, “Formation and interaction of membrane tubes,” *Physical review letters*, vol. 88, no. 23, p. 238101, 2002.

25. F. Hochmuth, J.-Y. Shao, J. Dai, and M. P. Sheetz, “Deformation and flow of membrane into tethers extracted from neuronal growth cones,” *Biophysical journal*, vol. 70, no. 1, pp. 358–369, 1996.

26. T. R. Powers, G. Huber, and R. E. Goldstein, “Fluid-membrane tethers: minimal surfaces and elastic boundary layers,” *Physical Review E*, vol. 65, no. 4, p. 041901, 2002.
